# Conditional termination of transcription is shaped by Rho and translated uORFS in *Mycobacterium tuberculosis*

**DOI:** 10.1101/2022.06.01.494293

**Authors:** Alexandre D’Halluin, Peter Polgar, Terry Kipkorir, Zaynah Patel, Teresa Cortes, Kristine B. Arnvig

**Author notes:** Institut de Biologie Physico-Chimique, CNRS UMR8261, Université de Paris, 75005 Paris, France. These authors contributed equally (Alexandre D’Halluin, Peter Polgar).

## Abstract

Little is known about the decisions behind transcription elongation *versus* termination in the human pathogen *Mycobacterium tuberculosis*. By applying Term-seq to *M. tuberculosis* we found that the majority of transcription termination is premature and associated with translated regions, i.e. within previously annotated or newly identified open reading frames. Computational predictions and Term-seq analysis upon depletion of termination factor Rho suggests that Rho-dependent transcription termination dominates all TTS including those associated with regulatory 5’ leaders. Moreover, our results suggest that tightly coupled translation, in the form of overlapping stop and start codons, may suppress Rho-dependent termination. This study provides detailed insights into novel *M. tuberculosis cis*-regulatory elements, where Rho-dependent, conditional termination of transcription and translational coupling together play major roles in gene expression control. Our findings contribute to a deeper understanding of the fundamental regulatory mechanisms that enable *M. tuberculosis* adaptation to the host environment offering novel potential points of intervention.

## INTRODUCTION

The control of gene expression plays a critical role in the pathogenesis of *Mycobacterium tuberculosis*, the cause of tuberculosis (TB) e.g.^1–4^. Transcription initiation is the first step of gene expression, and while the process of transcription initiation and the role of transcription factors are relatively well characterised in *M. tuberculosis*, reviewed in^5^, the role and the molecular mechanisms underlying the decision between transcription elongation and termination are poorly understood. Few canonical intrinsic terminators (ITs), i.e., a stem-loop followed by a poly-U tract (‘L-shaped’) have been identified, let alone investigated in *M. tuberculosis* and results are conflicting. The search for alternative mechanisms behind *M. tuberculosis* intrinsic or factor-independent termination has led to the suggestion that ‘I-shaped’ terminators, i.e. a stem-loop without poly-U tail may be sufficient^6^, but more recent findings indicate that, as a minimum, a partial or interrupted poly-U stretch needs to be in place for unidirectional, factor-independent termination to take place^7–10^.

A second mechanism of factor-independent transcription termination involves head-on collisions between transcription elongation complexes on converging genes. This is often associated with hairpins and accounts for a significant proportion of termination events in *Escherichia coli*^11^. Previous results suggest that converging genes in M. tuberculosis have a substantial antisense element, which could be associated with such a mechanism^7^.

A third mechanism of transcription termination is mediated by the hexameric helicase, Rho^12^. Rho binds to pyrimidine-rich stretches (*RUT* sites) on nascent RNA and induces transcription termination as much as 100 nucleotides downstream^12,13^. This, in combination with rapid post-termination 3’ processing has made identification of actual *RUT* sites challenging^14^. However, recent cryo-EM structure studies have provided additional length- and sequence specificity for the primary binding site on nascent transcripts in *E. coli*^15^.

*M. tuberculosis rho* is essential, suggesting that Rho-dependent (RD) termination of transcription plays an important role in the control of gene expression in this pathogen^16^. Several investigations into the function of Rho in other bacteria have employed the antibiotic bicyclomycin, which is generally a potent inhibitor of Rho function^12,14,17^. However, the binding of bicyclomycin to *M. tuberculosis* Rho is obstructed due to a leucine to methionine substitution, which also affects the translocation and termination efficiencies of the protein^18^. Using a Rho depletion strain (RhoDUC), in which the addition of anhydrotetracyclin (ATc) simultaneously represses transcription of *rho* and induces degradation of Rho protein, Botella *et al*. demonstrated that induction with ATc leads to a 50% reduction in the levels of Rho protein after six hours followed by cell death after 24 hours, highlighting the importance of RD termination of transcription in *M. tuberculosis*^16^.

Correct termination of transcription at the 3’ end of genes or operons is an integral part of the transcription cycle, while premature or conditional termination of transcription in the 5’ untranslated region (UTR) upstream of or early within an open reading frame (ORF), will suppress the expression of the downstream gene(s). Conditional termination of transcription is likely guided by specific physiological signals or ligands that modulate the potential of a terminator to promote transcriptional readthrough or termination. This mechanism, seen for example in the context of riboswitches and certain antibiotic resistance genes, offers an additional mechanism of gene expression control and potential intervention^19–23^. The application of Term-seq to map RNA 3’ ends has revealed an unexpected abundance of conditional intrinsic and RD terminators in other bacterial systems^17,21^. The decision between transcriptional termination and readthrough may be regulated by a variety of mechanisms including binding of metabolites, sRNAs or proteins to the transcripts^24–28^. Alternatively, the translation of small upstream ORFs (uORFs) and/or leader peptides may control transcription termination downstream (i.e. transcription attenuation), a mechanism that has received renewed attention in recent years as more potentially translated uORFs are being identified^17,23,29–32^. However, a global view of transcription termination in *M. tuberculosis*, including conditional termination is still lacking^33,34^.

To better understand fundamental aspects of *M. tuberculosis* post-transcriptional gene expression control and to identify conditional terminators associated with known and new 5’ leaders and/or riboswitches, we applied a combination of Term-seq^21^ and tagRNA-seq^35^ to map *M. tuberculosis* RNA 3’ ends. As expected, we found that few *M. tuberculosis* transcription termination sites (TTS) were associated with canonical ITs, while computational predictions combined with Term-seq and RNA-seq applied to the *M. tuberculosis* RhoDUC strain suggested that a large proportion of transcription termination events are likely RD. Moreover, most of the mapped TTS were associated with premature termination of transcription, some of which are likely associated with post-transcriptional control in response to specific molecular signals. Finally, by comparing TTS with ribosome footprints^36^, we found that premature termination is often associated with translation of small upstream ORFs (uORFs), located in regions previously considered as 5’ UTRs and sometimes overlapping with annotated genes, suggesting that translating ribosomes may contribute to the regulation of premature termination of transcription. Our study provides detailed insights into post-transcriptional control of gene expression in *M. tuberculosis*, with emphasis on conditional, RD termination of transcription, which appears to be counteracted by translational coupling. Moreover, it comprises a rich catalogue of novel *cis*-regulatory RNA leaders for further molecular investigations.

## RESULTS

### TTS identification and classification in *Mycobacterium tuberculosis*

To identify *M. tuberculosis* TTS on a genome-wide scale, RNA was isolated from triplicate, log-phase cultures of *M. tuberculosis* H37Rv, using previously published methods^7,37^; the RNA was subsequently analysed using Term-seq and tagRNA-seq to simultaneously map RNA 3’ ends and 5’ tri- and mono-phosphates, respectively, Figure 1^21,35^. To include the maximum number of transcription start sites (TSS) used in our analyses, we combined previously identified TSS^38^ with new TSS, which we obtained with tagRNA-seq (Table S1).

**Figure 1.**
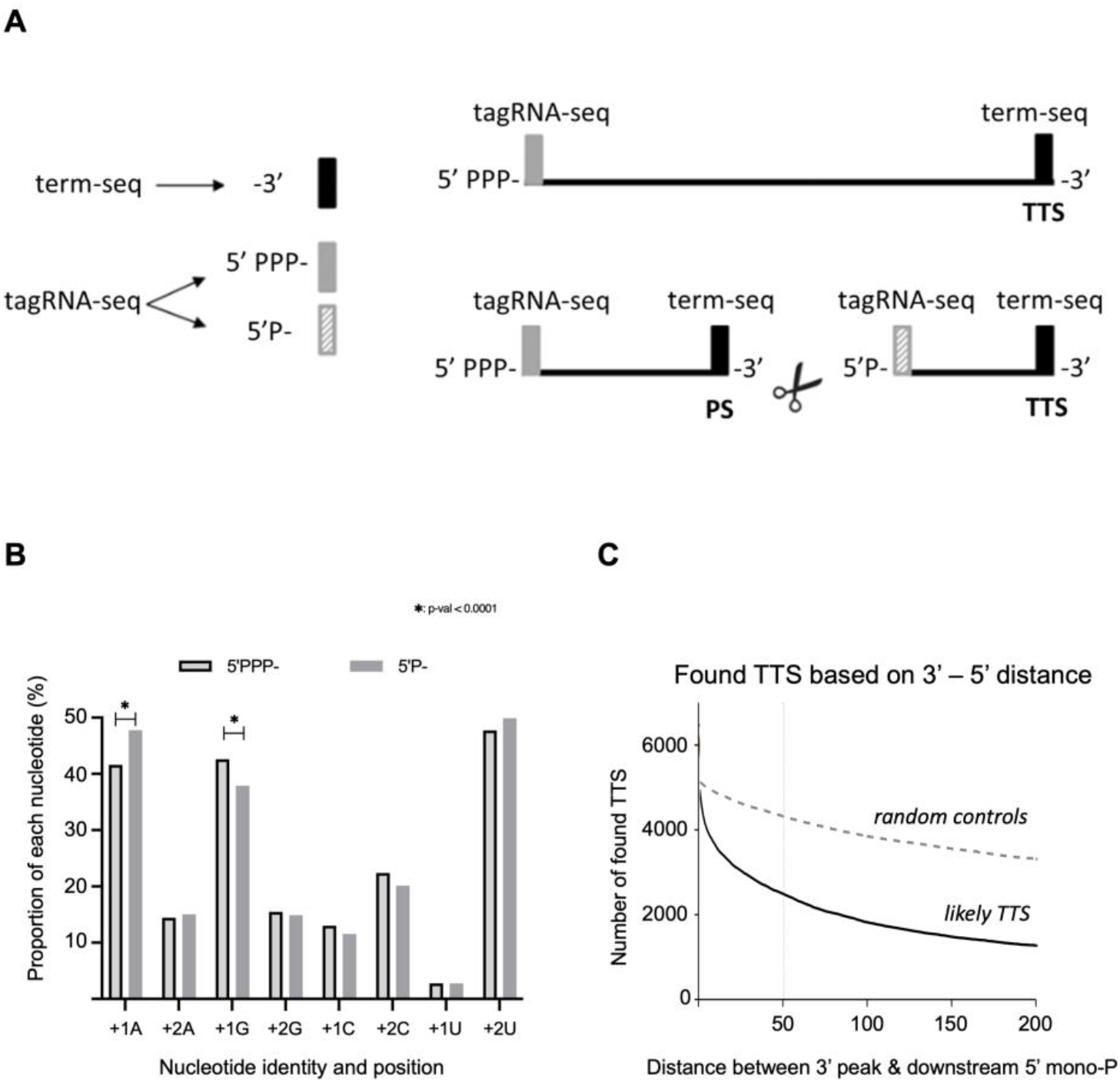
Transcription Termination Sites and Processing Sites. **A**. Termination of transcription generates a 3’ end, while internal cleavage of RNA will generate new 3’ and 5’ ends, in most cases 3’ OH and 5’ monophosphate, as shown in the bottom right panel. Term-seq enables the identification of RNA 3’ ends (3’, Black bar), while tagRNA-seq enables the distinction between tri-phosphorylated (5’ PPP-, grey bar) and mono-phosphorylated (5’ P-, hatched bar) transcripts. A 3’ signal followed by a 5’ P signal indicates a likely processed site (PS), while the remaining 3’ signals are classified as likely transcription termination sites (TTS). **B**. The proportion of each nucleotide within the first two positions was extracted and hypergeometric testing with FDR correction revealed an enrichment for transcripts with Adenine at +1 (15%, p-value =2,60137E-14) and an underrepresentation of Guanine at the same position (12%, p-value 7,10836E-09). There were no significant differences associated with position +2. **C**. The number of likely TTS, i.e. 3’ termini that are not accompanied by a 5’ mono-phosphate downstream was determined as a function of the 3’-5’ distance (black curve); grey, dashed curve shows randomly chosen positions plotted as a function of distance to downstream 5’ mono-phosphate.

Initial Term-seq reads indicated multiple shallow peaks throughout the transcriptome, which we attributed to imprecise termination and/or poor RNA polymerase (RNAP) processivity. By applying a cut-off of 4.8 counts per million (CPM, see methods), we obtained 5168 discrete, well-identifiable peaks throughout.

RNA 3’ ends identified by Term-seq may be generated by transcription termination or by endonucleolytic cleavage as part of RNA processing. Endonucleolytic cleavage will in most cases generate a 3’ OH followed by a 5’ monophosphate, the latter which can be distinguished from 5’ triphosphates resulting from transcription initiation at TSS by using tagRNA-seq ^35^ (Figure 1A). Subsequently, these monophosphates can be used to exclude potentially processed 3’ ends (processing sites, PS) and hence identify likely TTS. A minimum threshold of 1.27 CPM was determined for such likely PS (methods) resulting in 57,755 likely PS (Table S2). Of these, 1,235 (2.1%) coincided with the occurrence of 5’-triphosphorylated RNA ends identified as TSS. This suggests that these 5’ monophosphates result from tri-phosphate to mono-phosphate conversions rather than endonucleolytic cleavage. The *Bacillus subtilis* Nudix hydrolase, RppH, performing this conversion has a strict preference for a G-residue at the +2 position with slightly less stringent requirements for positions +1 and +3^39,40^. This is markedly different from the *E. coli* RppH, which show very little sequence preference^41^. To determine if there was any sequence bias in *M. tuberculosis* conversions, we extracted the sequence of the two first nucleotides associated with the 1,836 TSS (i.e. 5’ tri-phosphates) and from the 1,235 overlapping PS (i.e. 5’ monophosphates) and found that tri- to mono-phosphate conversions were enriched in transcripts that had A residues in the first position, while G residues underrepresented. Notably, we observed no significant differences associated with the +2 position (Figure 1B). Whether this translates into actual preference of the enzyme(s) involved requires further experiments. Next, we compared all RNA 3’ ends (obtained with Term-seq) to 5’ monophosphates (obtained with tagRNA-seq) located between 0 and 200 nucleotides downstream of the 3’ end in order to remove PS from the Term-seq data set and obtain TTS. Figure 1C shows how the number of likely TTS decreased as the 3’-5’ distance increased. We settled on an arbitrary distance of 50 nucleotides as our cut-off, identifying 2,567 likely TTS of the initial 5168 Term-seq peaks in the *M. tuberculosis* transcriptome (Table S3). These TTS separated into four main profiles shown in Figure 2: a single peak per gene/operon (e.g. *thiO*), multiple, discrete peaks within a defined region (e.g. *glyA2*), or distributed across entire genes and/or operons (e.g. *rpsJ/S10 operon*) and converging/overlapping sense and antisense peaks (e.g. *nrdB*). The latter profile is likely the result of RNAP collisions, recently reported to be abundant in *E. coli*^11^. Additional profiles can be seen in Figure S1.

**Figure 2:**
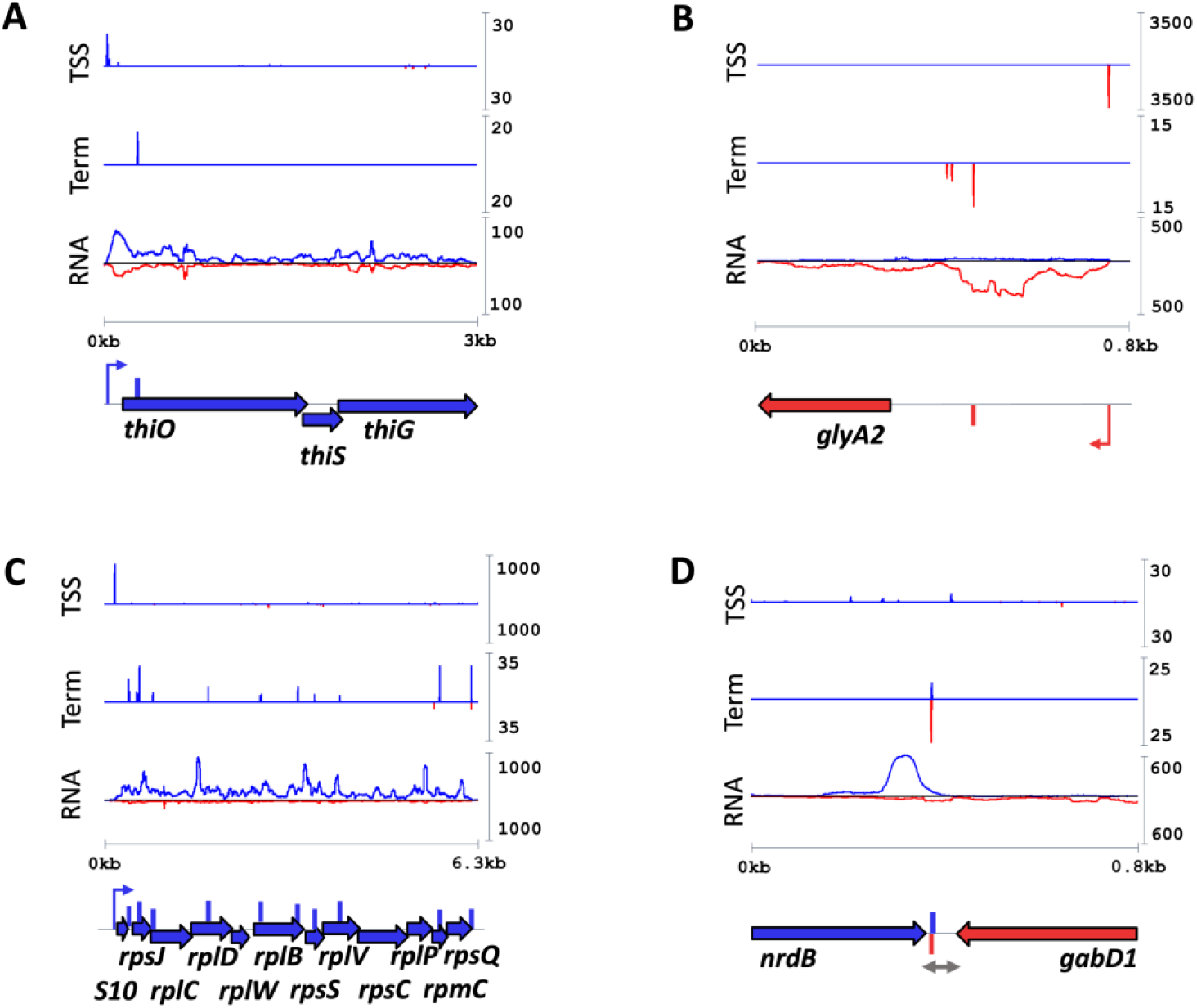
Transcription termination site (TTS) profiles. TTS profiles followed four main patterns: a single, sharp peak within a gene/operon (**A**); a cluster of peaks within a defined region (**B**), multiple, discrete peaks covering entire genes or operons (**C**) and converging peaks (**D**). Each plot shows one representative region with transcription start site (TTS) in the top panel, TTS profiles in middle panel and RNA-seq traces, bottom panel. Blue traces: Coverage on plus strand. Red traces: Coverage on minus strand. Horizontal Blue/Red arrows: ORF. Vertical Blue/Red bars: Mapped (dominant) transcription termination site (TTS). Hooked arrows: Mapped TSS.

All TTS were classified according to their position relative to nearest TSS upstream and annotated ORFs, using the definitions from^17^ with minor modifications (Figure 3). Thus, Internal TTS are located within annotated ORFs; Antisense TTS are antisense to annotated ORFs and their 5’ UTR (from stop Codon to TSS); Final is the dominant TTS located within 500 nucleotides downstream of the last ORF, (unless the region harboured another TSS or annotated gene); Orphans are TTS located in intergenic regions, including 5’ UTRs.

**Figure 3:**
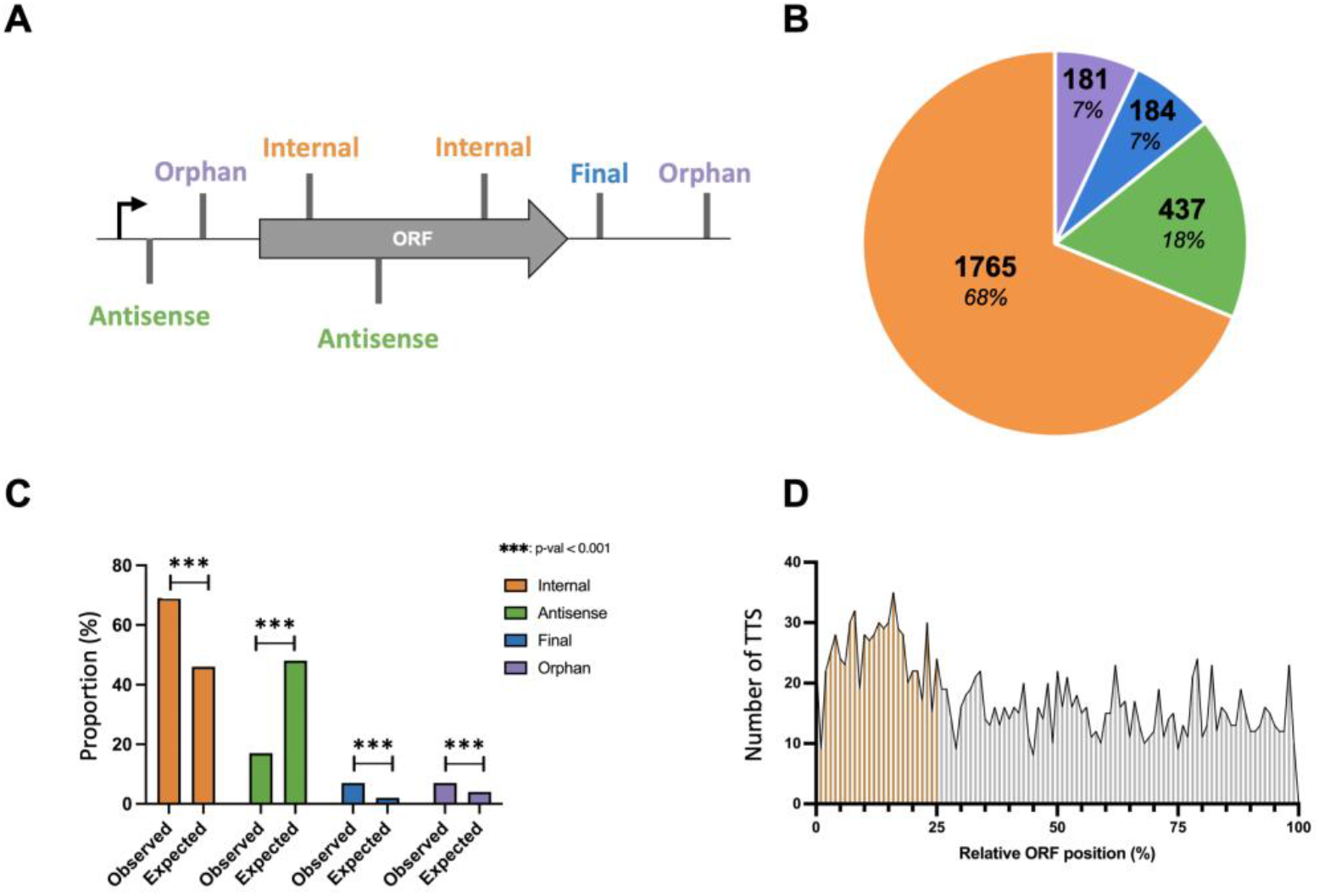
TTS classification and distribution. **A.** TTS were classified according to nearest upstream TSS and annotated ORFs. Grey vertical bars indicate TTS. Black vertical arrow: TSS. **B.** The number and proportion of each TTS class defined in **A**. **C.** indicates the proportion of observed TTS for each class versus expected TTS based on the extent of the regions within the genome; ***p-value<0.001 (Chi-square test with BH FDR correction). **D.** The distribution of Internal TTS across average ORFs.

According to these definitions, Orphan and Final TTS represented the smallest fraction at only 7%, followed by Antisense at 18%. The remaining two thirds of TTS were Internal, in other words, associated with translated regions. Comparing these fractions to the fraction that these categories represent within the genome revealed a significant enrichment of Internal (Chi-square FDR corrected, p-value = 1.46e-116), Final (p-value = 4.67e-76) and Orphan TTS (p-value = 2.44e-19), while Antisense TTS were significantly underrepresented (p-value = 5.64e-217; Figure 3C). Moreover, by mapping each Internal TTS relative to its cognate ORF size, we observed a further enrichment of TTS within the first quarter of ORFs. Together the results suggest that a large proportion of transcription termination in *M. tuberculosis* is premature and potentially controlled by specific *cis*-regulatory elements.

We have previously reported that *M. tuberculosis* 3’ UTRs can be several hundred nucleotides in length^7^. To determine the distribution of 3’ UTR lengths based on Term-seq data, we calculated the distance from the stop codon to nearest Final TTS downstream and plotted the frequency of each length (Figure S2). The results suggest that *M. tuberculosis* tends to have longer 3’ UTRs than e.g. *B. subtilis*^42^, offering the potential for post-transcriptional regulation.

### Intrinsic terminators are rare in *Mycobacterium tuberculosis*

*M. tuberculosis* is known for its paucity in canonical ITs^7,8,33^. Instead, alternative motifs and/or mechanisms of termination such as non-canonical ITs with imperfect or no poly-U tails or RD terminators have been suggested to play a role^6,8,9,16^. To gauge the relative contribution of individual mechanisms to transcription termination in *M. tuberculosis*, we compared our mapped TTS with different types of predicted terminators.

TransTermHP^43^, WebGeSTer DB^44^ and RNIE^9^ were used to predict ‘L-shaped’ ITs with perfect as well as imperfect U-tails, ‘I-shaped’ and ‘V-shaped’ ITs or Tuberculosis Rho-independent Terminators (TRITs), respectively. The resulting predictions all showed minimal overlap with mapped *M. tuberculosis* TTS, suggesting they each represent a small fraction of *M. tuberculosis* terminators (2%-4%; Figure S3A). Since these types of terminators are associated with a high degree of secondary structure, we used RNAfold^45^ to calculate the folding energy around each TTS and compared this to folding energies around random positions in the genome. As the plots were almost overlapping, we conclude that the mapped TTS were associated with limited secondary structure therefore corroborating the absence of structure-dependent terminators (Figure S3B). The most prominent overlap between prediction and experimental data was found with RhoTermPredict (RTP), which uses a two-step approach to predict RD terminators^13^. The first step identifies a 78-nt *RUT* site based on C:G ratio and regularly spaced C residues, while the second step searches for potential RNAP pause sites within 150 nucleotides downstream of the *RUT* site, which led to the identification of 23930 RD termination sites (RDTS) in *E. coli*^13^. We applied RTP with default parameters to the *M. tuberculosis* and *B. subtilis* genomes and found 29096 and 16390 predicted RDTS, respectively (Table 1; all predicted RTP sites for *M. tuberculosis* can be found in Table S3).

**Table 1.**

| Species | Genome size (Mbp) | % GC | Predicted RDTs | freq (RDTs/Mbp) |
| --- | --- | --- | --- | --- |
| *M. tuberculosis* | 4.4 | 66 | 29096 | 6612.7 |
| *E. coli* | 4.6 | 51 | 23930* | 5202.2 |
| *B. subtilis* | 4.2 | 44 | 16390 | 3902.4 |

\*From <sup>13</sup>

This indicates that RD terminators (as defined by Di Salvo *et al*.)^13^ are likely to be more abundant in *M. tuberculosis* than in *E. coli* and *B. subtilis*, in line with the GC content of these genomes. Of the 2567 mapped TTS, a large majority (1,879 or 73%) overlapped with predicted RDTS. Hypergeometric testing indicated that this was highly significant (p-value < 0.004) and suggests that RD termination of transcription may be the principal mechanism in *M. tuberculosis*.

### Depletion of Rho affects most *M. tuberculosis* TTS

To validate the prediction that a large proportion of the experimentally mapped TTS were potentially RD, we investigated transcription termination after Rho depletion. Duplicate cultures of *M. tuberculosis* RhoDUC ^16^ (kindly provided by D. Schnappinger) were grown to mid-log phase before the addition of anhydrotetracyclin (ATc) to induce Rho depletion. RNA was subsequently isolated at times 0, 3, 4.5 and 6 hours. Term-seq was applied and for each timepoint, the coverage of the 2,567 TTS mapped in H37Rv was extracted in a window of 2 nucleotides either side of the TTS, the profiles of TTS in H37Rv and uninduced RhoDUC being reasonably similar (Figure S4). A TTS score was calculated as CPM-normalised coverage at T0 relative to CPM-normalised coverage for Tt for all 2567 TTS and the distribution of scores was plotted for each timepoint. The plot, shown in Figure 4A suggests that for each timepoint, the coverage of more than half of TTS decreased over time as Rho was depleted (TTS score >1).

**Figure 4.**
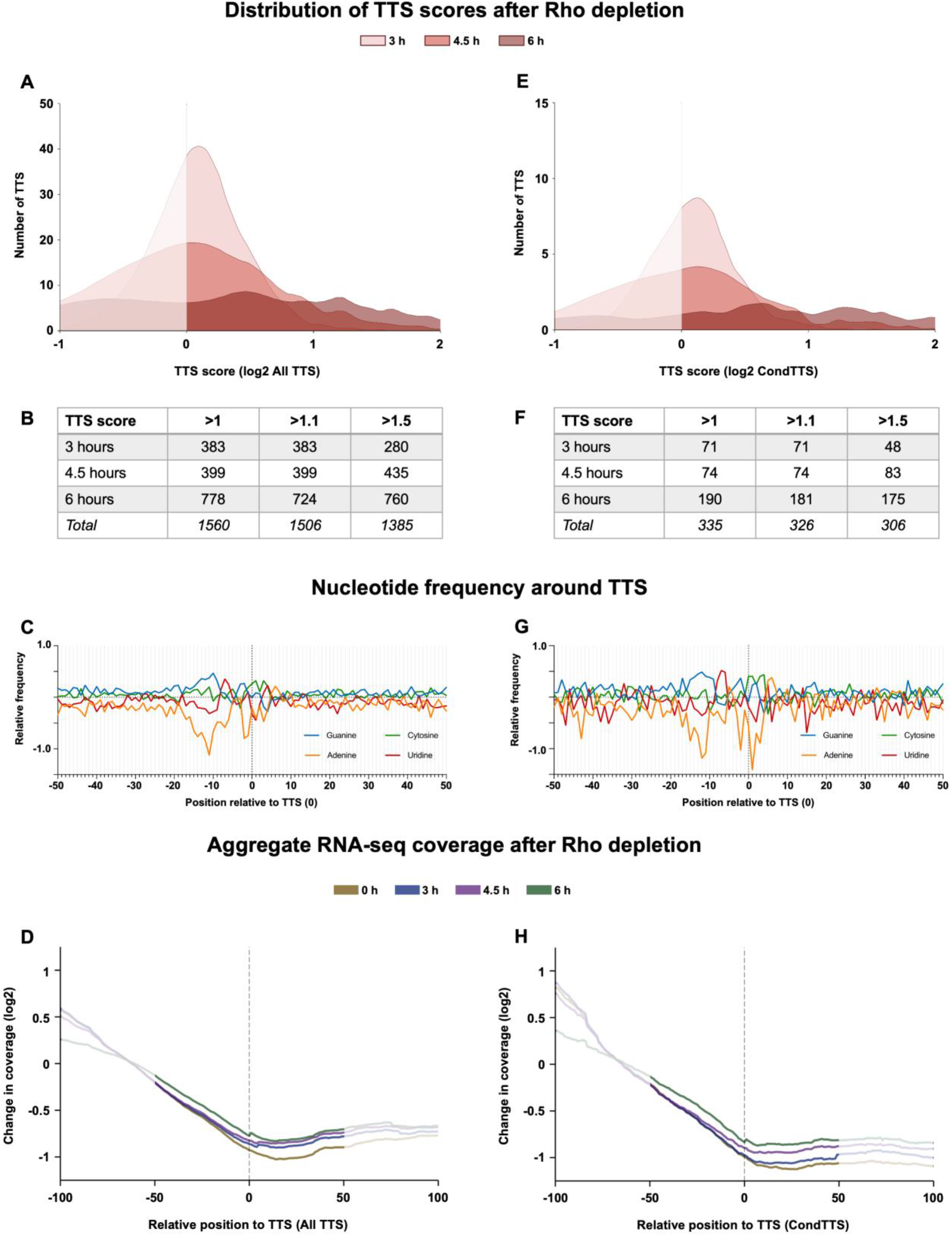
Changes in TTS coverage and readthrough following Rho-depletion. A TTS score (coverage at T0/coverage at T) was calculated for each H37Rv-mapped TTS and the distribution of TTS scores for all TTS (**A**) and CondTTS (**D**) plotted across the time course; shaded area indicates TTS scores <1. Tables with number of TTS obtained between each timepoint after FDR correction at different cut-offs are shown below (**B** and **F**). **C** and **G** show nucleotide frequencies associated with 1385/306 FDR corrected TTS (total at six hours). **D** and **H** show aggregate plots of the normalised RNA-seq coverage around each TTS generated from heatmaps shown in Figure S6.

Panel B in Figure 4 indicates the numbers of TTS identified as potentially RD based on increasing TTS scores at each depletion timepoint, for which the significance was validated with an FDR corrected t-test between each replicate. Thus, we observed a total of 1385 TTS decreasing > 50% from T_0_ to T_6_ (TTS score > 1.5, padj <0.05). Conversely, 300 TTS displayed a TTS score < 50% (padj <0.05; Table S4). The relationship between cut-off, q-values and resulting RD TTS has been shown in Figure S5.

This gradual and relatively subtle change is likely due to the nature of Rho depletion leading to more noisy data than instantaneous drug-mediated inactivation of Rho or the deletion of the *rho* gene, neither of which cannot be achieved in *M. tuberculosis*. To identify putative motifs associated with a positive TTS score, we plotted the nucleotide frequencies around the TTS assigned as RD and observed a slight enrichment of GC residues upstream of the mapped TTS with a marked underrepresentation of A residues from around −20 (Figure 4C). Although less pronounced, the profile ~5-15 nucleotides upstream of the TTS is to some extent reminiscent of the profile associated with *E. coli* RD terminators, except for the depletion of G-residues close to the TTS; the profile downstream of the TTS also did not display a depletion of G-residues nor a pronounced enrichment of C-residues as that reported for *E. coli*^14^. To obtain a better understanding of *M. tuberculosis* RD termination, we also measured transcriptional readthrough (RT) across the TTS by RNA-seq.

An RT-score was calculated for each timepoint based on the ratio of normalised read coverage upstream and downstream of each TTS (Table S4). In a complementary approach, we visualised changes in transcription around TTS by plotting RNA-seq coverage in heatmaps for each timepoint (Figure S6). Aggregate plots summarising these heatmaps illustrate changes in RNA-seq coverage around the mapped TTS (Figure 4D). Like the TTS score, the results suggest a gradual increase in transcriptional readthrough as Rho is depleted over time, where the most prominent difference could be observed between T=0 and T=3 hours (gold curve *versus* blue curve), while the differences between the remaining timepoints were less pronounced. In addition, aggregate plots of RNA-seq coverage around TTS located within predicted RD regions were generated, and these suggested a slightly better time-dependent resolution (Figure S6). Finally, we investigated the overlap between TTS with reduced coverage and TTS with increased RT since, in theory, there should be a good overlap between these if both are RD. The results suggested that around a quarter (406) of TTS identified as RD by each method were also identified by the complementary method, and we consider these high-confidence RD TTS. Using Fisher’s exact test, we found this overlap to be statistically significant (p-value = 0.001336, Figure 5A).

**Figure 5.**
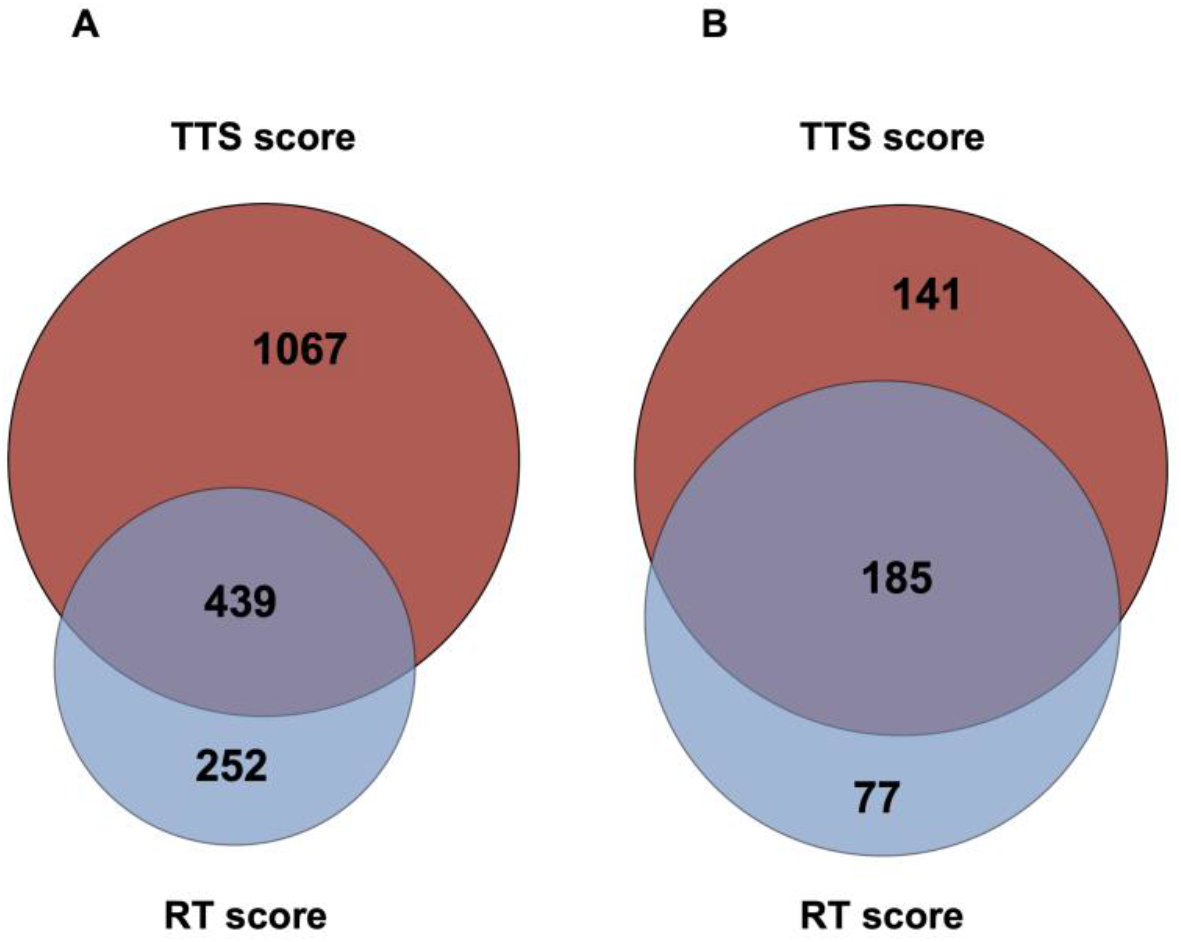
Overlaps between Rho-dependent TTS identified by different methods. Venn diagrams indicating the number of TTS identified as Rho-dependent (RD) after six hours’ depletion either by TTS score (red) or RT score (blue) and the overlap between these. The diagram in **A** shows that 29% of TTS classified as RD by TTS score were also classified as RD by RT score; conversely, 64% of RD TTS as determined by RT score overlapped with RD TTS as determined by TTS score (p-value = 0.001336, Fisher’s exact test). The diagram in **B** shows the same relations for CondTTS, where 52% of TTS classified as RD by TTS score were also RD by RT score, while 71% of RD TTS, as determined by RT score, were also classified as RD according to TTS score (p-value = 0.0009442, Fisher’s exact test).

### Conditional termination of transcription in *M. tuberculosis*

Our results show that >700 of Internal and Orphan TTS could essentially prevent transcription of full-length mRNA. These may represent conditional TTS often seen in the context of *cis*-regulatory elements such as RNA leaders and riboswitches^17,33,46^. To investigate conditional termination sites (CondTTS) associated with potential regulatory 5’ leaders, we focused on all Orphan TTS as well as Internal TTS within the first quarter of annotated ORFs, as we had found that this region was enriched for TTS (Figure 3D). Moreover, we only included the dominant TTS for each region. This led to the identification of 506 potentially regulated leaders, derived from primary TTS (123 Orphan TTS and 383 Internal), representing 20% of all TTS and potentially regulating more than 10% of annotated genes (Table S5).

As above, we plotted TTS scores and RT scores for all CondTTS (Table S5) over time (Figure 4D and F). We found that 60% of CondTTS were RD, seen as a statistically significant TTS score > 1.5 after 6 hours’ Rho depletion (Figure 4). The aggregate plot in Figure 4H indicates gradual time-dependent changes in RNA-seq coverage for CondTTS. Moreover, more than half of TTS identified as RD according to TTS score overlapped with those identified as RD according to increased RT (p-value = 0.0009442; Figure 5). Together these results suggest that RD termination of transcription may be more distinct for CondTTS than for the average *M. tuberculosis* termination event, although the nucleotide frequencies/patterns were similar in the two classes (Figure 4C and G)

Finally, we extracted the length of *M. tuberculosis* leaders from TSS to CondTTS, plotted these against their frequencies and analysed the distribution of longer RNA leaders (≥50 nucleotides in length) across different functional gene categories. The results suggested and uneven distribution with longer leaders enriched within Cell Wall & Cell Processes, Information Pathways and PE/PPE genes (p-value<0.05; Figure S7).

### *M. tuberculosis* CondTTS associated with translated uORFs

Small, translated ORFs are a prominent feature of bacterial genomes^17,23,29–32^. We noticed that several Orphan TTS fell within small to medium (≤100 nucleotides) ORFs located upstream of annotated ORFs; we refer to these upstream ORFS as uORFs. To gauge whether these uORFs were potentially translated, whereby Orphan TTS would become Internal TTS, we mined ribosome footprinting (Ribo-seq) data from Sawyer *et al*. to extract likely translated regions^36^.

*M. tuberculosis* has 1434 5’ leaders, defined as the region between mapped TSS and ORFs annotated in Mycobrowser release v4. Of these, 649 harboured Ribo-seq coverage of ≥250 reads (see methods), indicating these are translated leaders. Within these, we identified likely translated uORFs with the following characteristics: a minimum length of 5 AA starting with NUG (where N is any of the four nucleotides) and being either leaderless (89) or with a canonical Shine-Dalgarno (407, see methods), i.e. 596 likely translated uORFs associated with 383 5’ leaders (on average 1.5 uORF per leader region, Table S6). Orphan TTS were significantly enriched (2.2-fold) in translated readers (73 within 383, 19.1%) compared to 123 in all 1434 leaders (8.6%, p-value < 2.22e-16, hypergeometric test). This indicates that translation is intimately linked to the regulation of transcription termination, in agreement with a pervasive role of Rho.

The search also revealed that several of the identified uORFs overlapped by one or four nucleotides with either an annotated ORF or with another uORF downstream, suggesting that translation of the two ORFs may be coupled via a Termination-Reinitiation (TeRe) mechanism, possibly promoting tighter coordination of expression^47,48^. Since the majority of all TTS were Internal and CondTTS were significantly enriched in translated leaders, we hypothesized that translational coupling, similar to coupling of transcription and translation might impact transcription termination^12^. Hence, we decided to investigate the co-occurrence of RD TTS and overlapping ORFs.

First, we performed a systematic search for overlaps across the *M. tuberculosis* genome. The results revealed that 775 (19%) of annotated ORFs overlap with another annotated ORF upstream, indicating that ORF overlaps are widespread in *M. tuberculosis*, as previously reported^47^. The most common configuration involved direct overlaps between stop and start codons, likely associated with a TeRe mechanism of translation^47^. Of these, four-nucleotide NUGA (where N denotes any nucleotide) overlaps were almost five-fold more abundant than one-nucleotide URRUG (where R denotes a purine) overlaps (515 or 66% *versus* 110 or 14%), while longer overlaps appeared to favour the +1 frame over the +2 frame (Figure 6). Notably, overlapping ORFs were particularly abundant in genes associated with virulence, detoxification and adaptation including several Toxin-Antitoxin (TA) modules; i.e *relBE* and *relJK*, 28 of 48 *vapBC* modules, 2 of 9 *mazEF* modules, both *parDE* modules and *higAC* had NUGA overlaps (Figure 6E and Table S7).

**Figure 6:**
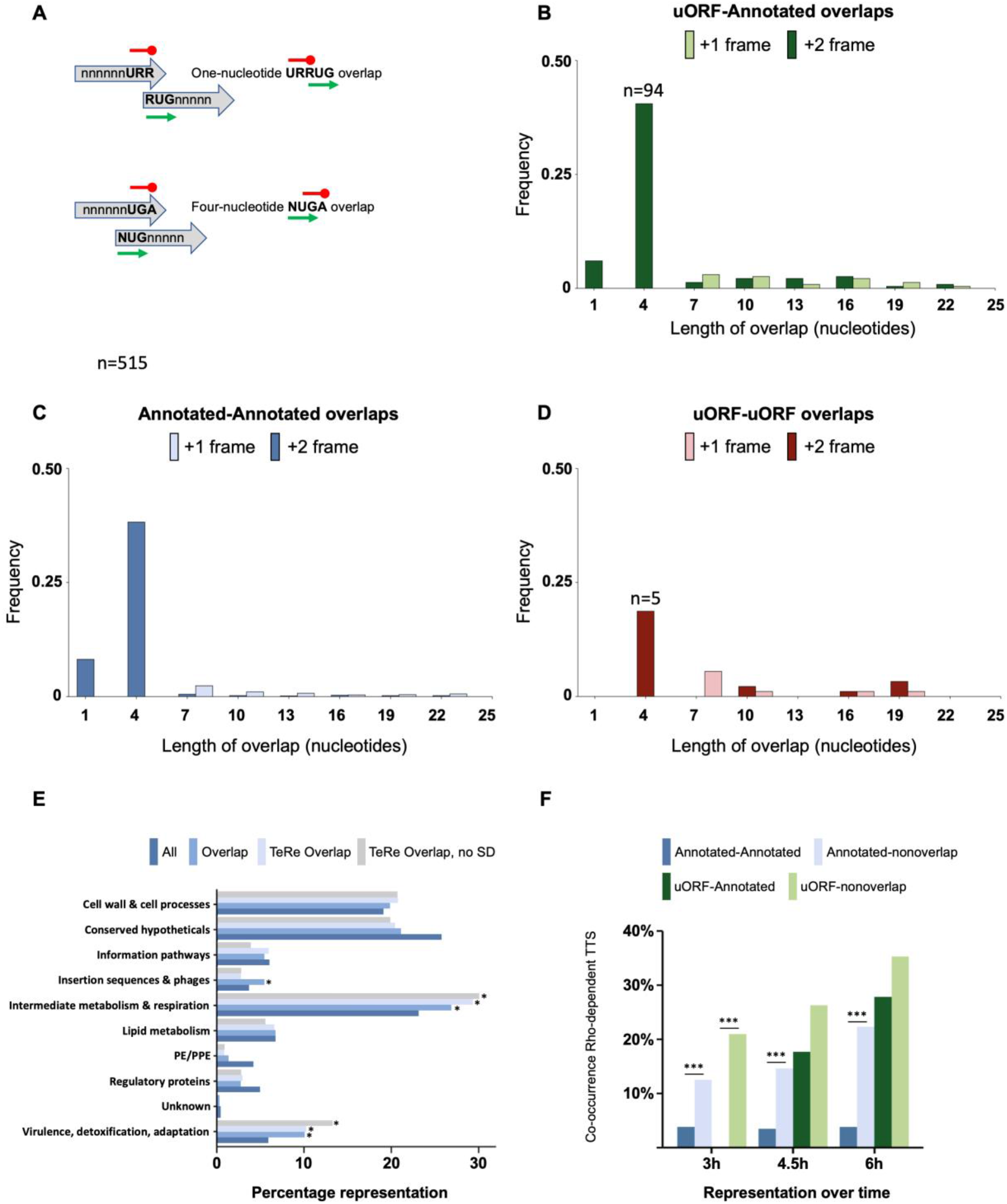
*M. tuberculosis* overlapping ORFs. A search for overlaps between all *M. tuberculosis* ORFs annotated in Mycobrowser^54^ indicated that 19% of all ORFs overlap between 1 and 25 nucleotides with an upstream ORF. The most abundant constellation was a four-nucleotide overlap, NUGA (N=any nucleotide) followed by a one-nucleotide overlap, URRUG (R=purine, with the restriction that a G has to be followed by an A), indicated in the schematic in panel **A**. Stop and start of the overlapping frames have been indicated in red and green, respectively in panel **A**. Panels **B-D** shows the breakdown into uORF-annotated gene overlaps, annotated-annotated and uORF-uORF overlaps, respectively; note that two- and five-nucleotide overlaps are not possible with conventional stop and start codons. Light and dark shaded bars indicate +1 and +2 frames relative to the upstream ORF, respectively. **E** shows the result of hypergeometric test of overlaps within functional gene categories (with BH corrections for FDR). **F** indicates co-occurrence of overlaps from **B**, **C** and **D** with TTS identified as RD by TTS score or RT score in Tables S4 and S5, with significance (p-value ≤ 0.05 Fischer test with BH corrections).

Applying the same search to overlaps between uORFs and annotated ORFs or uORF-uORF pairs showed the same trend, i.e. four-nucleotide and one-nucleotide overlaps being the most and second-most abundant type, respectively (Figure 6D and E, Table S7). This indicates that overlapping ORFs sharing the active site within a ribosome are a common feature in *M. tuberculosis*. Like all *M. tuberculosis* ORFs, the initiating nucleotide of the second ORF was dominated by purines with only a minor fraction of UUG and no CUG starts.

Notably, about 58% of ORFs initiated with a NUGA or URRUG overlap did not have a canonical SD (as defined above), although a weaker or non-canonical SD might be present. Finally, we calculated the co-occurrence of likely RD TTS and overlapping *versus* non-overlapping ORFs. Figure 6F summarises the results, which suggest that transcription within non-overlapping ORFs is significantly more likely to undergo RD termination than transcription within overlapping ORFs. This was most obvious at 3 hours and more so for uORF-annotated (dark green *versus* light green in Figure 6F) overlaps than for annotated-annotated (dark blue *versus* light blue, Figure 6F) overlaps, but also at 4.5 and 6 hours we observed a significant difference in RD termination between overlapping, annotated ORFs and their non-overlapping counterparts. Together with the finding that transcription termination was more abundant in translated regions, our results suggests that many uORFs are potentially regulatory by underpinning RD termination of transcription, while RD termination of transcription itself may be suppressed by a tighter coupling of translation (specifically 1- and 4-nt overlaps), which applied to all ORFs.

### Translation of *M. tuberculosis* uORFs and overlapping ORFs

Naturally, the presence of a ribosome footprint does not necessarily mean that a given uORF is translated. Therefore, to probe whether selected uORFs were translated, we made in-frame *lacZ* fusions and expressed these in *Mycobacterium smegmatis*. We selected leaderless (*rne*), isolated (*pe20, Rv1535, ilvB1*) and overlapping (*dnaA, glyA2, rpfA*) uORFs. The results demonstrated that all of the selected uORF-*lacZ* fusions are translated in *M. smegmatis* to different extents, suggesting that the uORFs are also translated in *M. tuberculosis* (Figure 7). This in turn supports that the associated CondTTS were Internal rather than Orphan.

**Figure 7:**
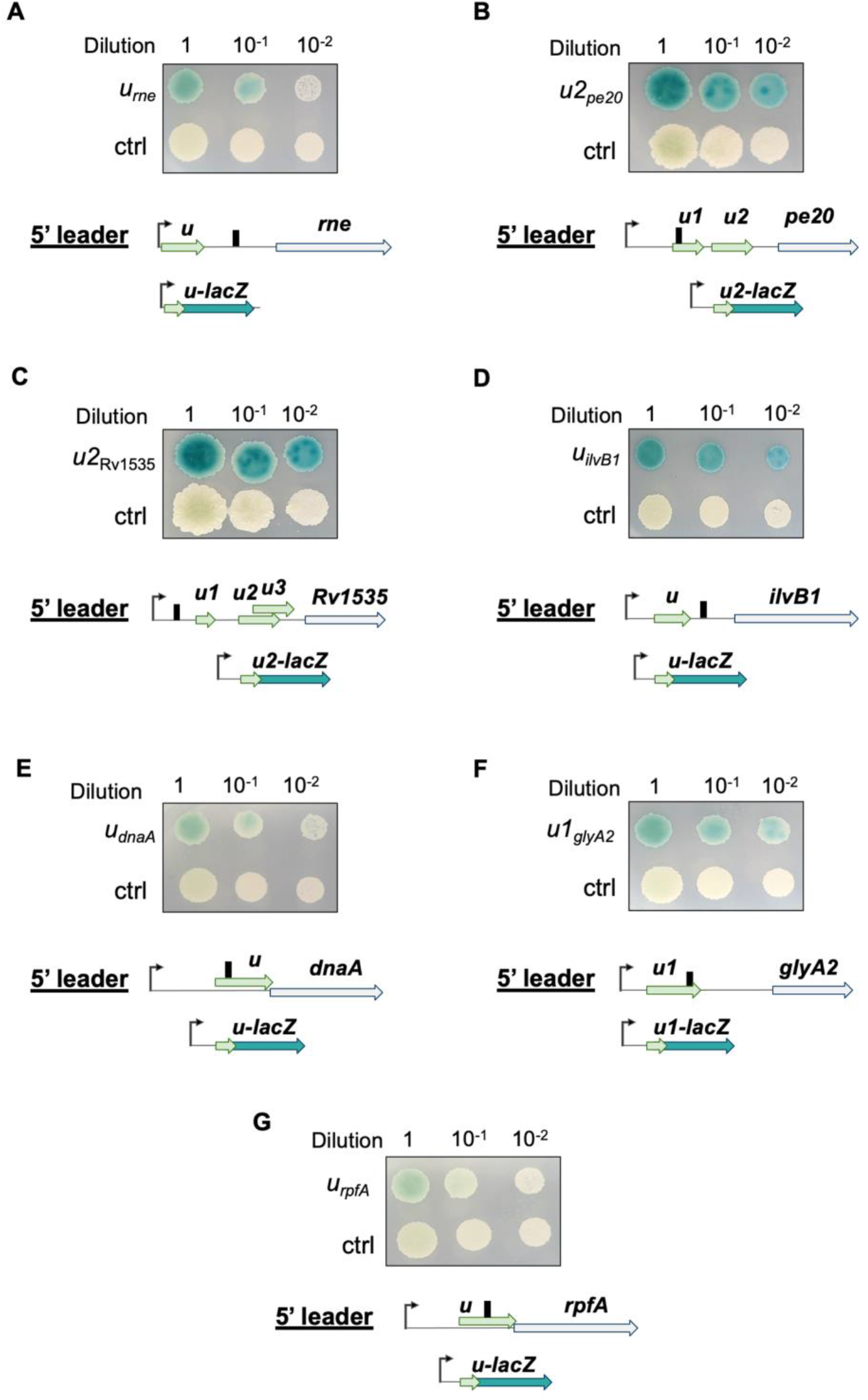
Translation of CondTTS-associated uORFs. Translation of TTS-associated uORFs was validated using in-frame *lacZ* fusions. Translation of selected uORF was validated by in-frame *lacZ* fusions of 20 nucleotides upstream of the identified SD to four codons into the uORF. Black arrow: TSS. Blue arrows: Annotated ORF. Light green arrow: Newly identified uORFs. Turqoise arrow: *lacZ*

Interestingly, some of the identified uORFs resided within the aptamers of known riboswitches including the Glycine, Cobalamin and YdaO riboswitch aptamers, respectively. We are currently investigating how translation affects riboswitch function and *vice versa*.

## DISCUSSION

The aim of this study was to define at nucleotide resolution, where in the *M. tuberculosis* genome, transcription termination happens and by which mechanism(s). To this end, we applied Term-seq to *M. tuberculosis* RNA and found indications for i) RD termination of transcription is the dominant mechanism across the genome; ii) premature termination of transcription is pervasive in *M. tuberculosis*; iii) conditional terminators are often associated with translated uORFs; iv) overlapping ORFs are abundant and potentially suppress RD transcription termination.

After applying a 4.8 read depth cut-off, we observed four main types of TTS profiles, which all suggested specific termination sites, seen as discrete peaks. However, before applying this cut-off, we observed several shallow, multi-peak profiles, suggesting inefficient and imprecise transcription termination, potentially related to the reduced translocation efficiency of *M. tuberculosis* Rho, which also results in multiple TTS *in vitro*^18^. Alternatively, *M. tuberculosis* RNAP could exhibit poor processivity as a result of a slower transcription elongation rate^49^. Further investigations are required to resolve this question.

Based on our analyses, we concluded that all TTS, but particularly CondTTS, often associated with specific *cis*-regulatory elements such as riboswitches or protein-binding leaders, were dominated by RD termination. Using TTS coverage, we found that a slightly higher proportion of CondTTS (60%) than all TTS (54%) displayed Rho dependence, identified by a statistically significant TTS score > 1.5 after 6 hours’ depletion and corroborated by a more extensive overlap of RD TTS identified by the different methods (Figure 5). Overall, our results suggest that the majority of *M. tuberculosis* TTS were dominated by RD termination, which is substantially more than reported for *E. coli* (>30%), although the study by Dar and Sorek referred to genes rather than TTS^14^. This indicates that, in line with predictions, *M. tuberculosis* resides at one end of a spectrum with terminators being almost exclusively RD, *B. subtilis* positioned at the opposite end with intrinsic terminators dominating and Rho being less important and *E. coli* in the middle, making ample use of both types of termination^33^.

Our results also suggest that even based on our stringent data analysis >10% of all genes are controlled by 5’ leaders with robust and specific CondTTS, seen as strong, discrete TTS peaks. This estimate is based on a (conservative) definition of leaders reaching no further than the first 25% of an ORF, the number may well be higher, and we did observe several CondTTS further into ORFs e.g. within multi-cistronic operons, where they likely regulate expression of the downstream ORF(s) potentially modulating the ratio of gene products within operons.

A further comparison between *M. tuberculosis* TTS locations to those of *E. coli* TTS^17^, highlights another major difference between the two species. Although Internal TTS (i.e. within ORFs) was by far the largest class in both organisms, we note a more than 2-fold difference in their relative abundance (68% in *M. tuberculosis versus* 29% in *E. coli*). At the same time, the TTS flanking either end of ORFs, i.e. Orphan (5’) and Final (3’) represented a much lower fraction in *M. tuberculosis* than in *E. coli* (<7% *versus* >20%). Together, this suggests that *M. tuberculosis* TTS tend to be more associated with coding regions than *E. coli* TTS. This could to some extent be ascribed to differences in ORF annotations and data analysis, but the fact that the proportion of Antisense TTS was almost identical in the two species indicates that the difference reflects actual, biological variations. The pervasive role of Rho combined with a lack of intrinsic terminators in *M. tuberculosis* likely contribute to this disparity, as RD termination is generally considered linked to translation. Moreover, the reduced motor efficiency of *M. tuberculosis* Rho^18^, could also play a role and potentially lead to some degree of readthrough from untranslated (Orphan) into coding (Internal) regions. Finally, although we did re-assign some Internal TTS to Final TTS, it is possible that post-termination 3’ trimming varies due to variation in 3’ structures and the complement of nucleases in the two species^14,50^.

Our results further indicated that many of the identified leaders harboured coding regions, in line with previous findings^30,31^, and furthermore suggested that CondTTS were more abundant in translated leaders. We did find some commonality (i.e. 65 identical and 87 isoforms) between our proposed translated uORFs and those recently reported by Smith *et al*.^31^. These differences are likely due to variations in experimental approaches as well as defining parameters such as for example peptide length. It remains unclear at this stage whether translated uORFs are regulatory or encoding peptides with *trans* functions or both, but at least in some cases, the levels of expression and amino acid conservation indicate functional peptides (manuscript in preparation).

We found that closely overlapping ORFs (i.e. by one and four nucleotides), including uORFs, are common in *M. tuberculosis* with four-nucleotide overlaps (+2 frame) being almost five-fold more abundant than one-nucleotide overlaps (+1 frame). This may to some extent be due to the sheer number of variations associated with each motif; i.e. URRUG is more or less limited in five positions, while NUGA(N) is only limited in three positions. At the moment, we cannot explain why longer overlaps seem to favour the +1 frame. Of note, more than half the downstream ORFs were devoid of canonical signals for translation initiation, suggesting tight translational coupling. We are currently investigating how such overlaps impact translation of SD-less downstream ORFs. It has been suggested that translational coupling can be via 70S scanning, which requires a SD^51^ or via TeRe, which may involve 30S as well as 70S ribosomes and not necessarily requiring an SD^47^. A quarter of *M. tuberculosis* ORFs are expressed from leaderless transcripts^38^ i.e. initiating translation with 70S ribosomes, indicating a propensity towards maintaining ribosomes in their assembled state. We therefore propose that the ‘handover’ from one ORF to a downstream ORF is likely to involve 70S ribosomes, which may be a means of saving energy in addition to coordinating gene expression and, according to our findings, suppress RD transcription termination.

Like transcriptional units or operons, *M. tuberculosis* appears to have translational units where two or even more ORFs overlap, and where expression of an ORF may to some degree depend on the translation of its upstream ORF (Kipkorir *et al*. manuscript in preparation). This seemingly non-canonical translational control is in line with previous findings of an extensive leaderless transcriptome and uORF-dependent gene expression^29–31,36,38^.

It remains to be established whether transcription and translation in *M. tuberculosis* are coupled, as in *E. coli*^52^ or mostly uncoupled as in *B. subtilis*^53^. On one hand, the principal role of Rho in transcription termination would suggest that coupling is likely in order to avoid unintended, premature termination. On the other hand, the proposed reduced motor function of *M. tuberculosis* Rho might reduce the need for coupling. Either way, the picture that emerges is that a significant proportion of *M. tuberculosis* gene expression control is post-transcriptional in the form of RD, conditional termination of transcription, translational coupling via TeRe or a combination of these. Although there is still much to be learnt, this study contributes our understanding of fundamental regulatory mechanisms in a pathogen that does not conform to the images of model organisms.

## LIMITATIONS OF THE STUDY

The main aim of this study was to identify on a genome-wide level transcription termination sites and the mechanisms behind these in *M. tuberculosis*. However, several studies have indicated that 3’ trimming of terminated transcripts is rapid and that the distinction between terminated and trimmed ends may be difficult without including nuclease mutants and/or in vitro transcription assays. As the current study was done using high-throughput sequencing methods on nuclease proficient strains, future studies employing such validations may be able to more accurately distinguish termination from 3’ trimming in *M. tuberculosis*. Moreover, the fact that *M. tuberculosis* Rho cannot be chemically inactivated by e.g. bicyclomycin means that the data will inevitably be more noisy as the depletion develops over several hours. Although some of our findings are based on statistically significant correlations, these require different more targeted approaches to validate the proposed mechanisms.

## ACKNOWLEDGEMENTS

The authors are grateful to Dirk Schnappinger, Weill Cornell for providing the RhoDUC strain, and to Fabian Blombach and Finn Werner, UCL for helpful suggestions and critical reading of the manuscript.

## AUTHOR CONTRIBUTIONS

Conceptualization, K.B.A.; Methodology, K.B.A., A.D., P.P., T.C.; Investigation, A.D., P.P, T.K., Z.P., Formal analysis, K.B.A., A.D., P.P.; Writing – original draft, K.B.A., A.D., P.P.; Writing – review and editing, K.B.A., A.D., P.P, T.K., Z.P, T.C.; Funding acquisition, K.B.A., T.K., T.C.

## STAR METHODS

### RESOURCE AVAILABILITY

Further information and requests for resources and reagents should be directed to and will be fulfilled by Kristine B. Arnvig

#### Materials availability

New plasmids generated in this study can be obtained by contacting the lead author

#### Data and code availability

All sequencing data files are available on ArrayExpress with the accession number E-MTAB-11753. The codes used to generate results and corresponding figures are all available at https://github.com/ppolg/Mtb_termseq.

### EXPERIMENTAL MODEL AND SUBJECT DETAILS

#### Bacterial strains and growth conditions

*M. tuberculosis* H37Rv and *Mycobacterium smegmatis* MC^2^ 155 were cultured on Middlebrook agar 7H11 supplemented with 10% OADC (Sigma), 0.5% Glycerol and 50 μg/ml hygromycin if appropriate and in liquid Middlebrook 7H9 supplemented with 10% ADC (Sigma), 0.5% Glycerol, 0.05% Tween 80 and 50 μg/ml hygromycin where appropriate. Cultures were harvested at an OD_600nm_ ~0.6 for mid-log phase. *M. tuberculosis* RhoDUC ^16^ was grown as previously described with 50 μg/ml hygromycin, 20 μg/ml kanamycin and 50 μg/ml zeocin. At OD_600nm_~0.6, depletion of Rho was induced by the addition of 500 ng/ml anhydrotetracyclin (ATc) and cells were harvested after 0, 1.5, 3, 4.5 and 6 hours. *Escherichia coli* DH5α was used for all cloning and were cultured on solid LB 1.5% agar or in liquid LB supplemented with 250 μg/ml hygromycin.

### METHOD DETAILS

#### Molecular cloning

pIRATE2020 was made by replacing the region in pIRATE^37^ between Xho I and Hind III with a region harbouring a shorter polylinker, slightly modified promoters driving the expression in both directions and an intrinsic terminator between the two promoters (See Table S9 for the inserted sequence). All subsequent pIRATE plasmids, listed in Table S8, were constructed using Gibson assemblies with oligos (Sigma) or geneBlocks (IDT) listed in Table S9. Generally, inserts spanned the region from 20 basepairs upstream of the presumed SD sequence and 20 basepairs upstream to the first four-six amino acids of the tested uORF. Once plasmids with the desired sequence had been identified, these were transformed into *M. smegmatis* by electroporation.

#### Translational reporter gene fusions

In-frame *lacZ* fusions were made in pIRATE2020 (Tables S8 and S9) by cloning inserts between the Hind III and Nco I sites; all were expressed in *M. smegmatis*. Spotting assays were done with 5 μl *M. smegmatis* at OD_600_~0.6 at the dilutions indicated.

#### RNA isolation and purification

*M. tuberculosis* and *M. smegmatis* RNA was extracted as previously described^7,55^. Briefly, cells were cold shocked by adding 30% ice directly to cultures, followed by centrifugation at 5000 rpm for 10 minutes at 4°C and finally RNA extraction using the FastRNA Pro Blue Kit (MP Biomedicals) following the instruction of the supplier. RNA concentration and purity was assessed using Nanodrop 2000 (ThermoFisher). Residual genomic DNA was removed using Turbo DNase (ThermoFisher) according to manufacturer’s instructions, followed by extraction with phenol-chloroform and ethanol precipitation. The RNA was subsequently checked for DNA by PCR using RedTaq readymix (Sigma). RNA integrity was assessed using 2100 Bioanalyzer (Agilent) before library preparation and RNA sequencing.

#### Library preparation and RNA sequencing

Library construction and sequencing were handled by Vertis Biotechnology AD (https://www.vertis-biotech.com/home), where all details are available.

Libraries were sequenced on an Illumina NextSeq 500 using 1×75 basepair read length. RNA sequencing quality control was assessed using FastQC. Sequences were mapped to the genome of *M. tuberculosis* AL123456.3 using Bowtie2^56^ and reads mapping more than once were discarded using Samtools^57^. Coverage was extracted using Bedtools^58^. For each dataset, the coverage was normalised to counts per million. All RNA-seq traces were visualised using Artemis^59^.

#### Transcription Start Site (TSS) and Processed Site (PS) mapping

Three biological replicates of *M. tuberculosis* H37Rv were harvested at OD_600nm_~0.6, and total RNA sequenced with tagRNA-seq^35^. The TSS and PS were extracted as below.

For TSS mapping, the coverage of the 5’ end of reads was extracted using Bedtools for 5’ tri-phosphate RNA fractions. The coverage at each position was normalised to counts per million and a geometric mean calculated from the three biological replicates. A threshold of 50 normalised reads was used to extract peaks at a single nucleotide level and were merged in a window of 3 nucleotides. Each TSS was with respect to the first downstream feature in a window of 500 nucleotides and compared to data from Cortes *et al*.^38^. This resulted in 59 new TSS that were added to the data from Cortes *et al*. and used in the screening of RNA leaders described below.

For PS mapping, coverage at the 5’ end of reads from the 5’ mono-phosphate RNA fraction was extracted using Bedtools and normalised to counts per million. The geometric mean across all three replicates was calculated at every position. Peaks were filtered against a background threshold, calculated as 1.27 CPM based on the mean coverage in intergenic regions. This resulted in the identification of 57,755 PS, which were used to filter termination sites, as described below.

#### Transcription Termination Site (TTS) mapping

*M. tuberculosis* RNA used for tagRNA-seq were also sequenced by Term-seq^21^ and transcription TTS extracted as outlined below.

Read coverage was extracted using Bedtools, normalised to counts per million, and the geometric mean calculated for each genomic position across all three replicates. A defined threshold was calculated using a similar approach as that applied to PS, described above. Within the same ORF, a TTS was assumed to be primarily associated with either the start (for conditional termination) or end (for RNA 3’ trimming) of the ORF, (https://github.com/ppolg/Mtb_termseq/blob/main/R/Mtb_peaksfrequency.R). Based on this rationale, each annotated ORF was divided in three equal length and the middle third of each ORF used to screen for background coverage levels. Following this analysis, a median read depth of 4.80 CPM was used as background cut-off for TTS mapping.

Bedgraph files were generated using deeptools^60^ on both strands for each biological replicate. Term-seq peaks based on this data was extracted using the package termseq_peaks^17^, with default parameters. Peaks located within a 100-nucleotide window were filtered and only the dominant peak, identified as the highest CPM peak was mapped.

We checked the presence of PS sites within a 1- to 200-nucleotide windows (see Figure 1C; https://github.com/ppolg/Mtb_termseq/blob/main/R/Mtb_PS_distance.R) filtering in a 50-nucleotide window downstream of the newly identified 3’ends. The 2567 Term-seq) downstream of the newly identified 3’ends. The 2567 Term-seq peaks without a processing site within this downstream window were identified as a TTS.

TTS were classified according to their genome position and localisation compared to features using Bedtools; TTS within an annotated ORF, antisense of a gene (ranging from its TSS or start codon to the stop codon), and the strongest TTS located within 500 nucleotides downstream of a gene was classified as Internal, Antisense and Final, respectively. TTS not falling into either of the three categories above (due to being intergenic or located between a TTS and the start codon) were classified as Orphan. (Figure 3A).

#### Computational prediction of transcriptional terminators

Experimentally identified TTS genomic positions were compared to predicted terminators using software based on different parameters. TransTermHP (http://transterm.cbcb.umd.edu) predicts L-shaped intrinsic terminators, based on a stem-loop, flanked by an upstream A-tail and a downstream T-tail within 15 nucleotides^43^. WebGeSTer DB (http://pallab.serc.iisc.ernet.in/gester/index.html) predicts L-, I-, U-, X- and V-shaped intrinsic terminators based on the best stable structure downstream of a stop codon^44^. RNIE is based on covariance using the Infernal package, which allows the discovery of the only conserved motif of terminators in *M. tuberculosis*, TRIT (Tuberculosis Rho-Independent Terminator)^9^. RhoTermPredict (RTP) searches for consensus *RUT* sites with a high C:G ratio followed by a palindromic sequence (stemloop) for RNA polymerase pausing^13^. RTP was run with default parameters and subsequently with reduced downstream window as described with all predictions listed in Table S3.

The free folding energy of a region of 40 nucleotides upstream and 10 nucleotides downstream of the TTS was calculated using RNAfold^45^ on sequences extracted using Bedtools.

#### TTS score and Readthrough score after Rho depletion

Two biological replicates of *M. tuberculosis* RhoDUC strain were harvested before and 3, 4.5 and 6 hours after the onset of Rho-depletion. Total RNA was sequenced with RNA-seq and Term-seq. Based on the Term-seq alignment, the 3’ end coverage was extracted for each biological replicate, normalised to counts per million (CPM) and the geometric mean calculated at each genomic position. Read coverage of the TTS was extracted in a window of 5 nucleotides (2 nucleotides on either side of the extracted TTS position from H37Rv), and the values compared between the *M. tuberculosis* H37Rv and RhoDUC strains. The log_2_ variation between RhoDUC and H37Rv was calculated, and MA plotted (Figure S4).

TTS scores at times 3, 4.5 and 6 hours were calculated by comparing coverage in these 5-nucleotide windows between time 0 and each subsequent timepoint. TTS scores with a value >1.1 and an adjusted p-value <0.05 (based on a t-test with Benjamini-Hochberg FDR corrections) were considered significantly decreasing.

A readthrough score (RT-score) was calculated for each timepoint from RNA-seq data in a similar fashion, by comparing coverage in windows of 100 nucleotides upstream (US) and downstream (DS) of each identified TTS calculated by HTseq-counts with the “union” parameter^17,61^. RT was calculated as a ratio of DS and US. RT-scores were calculated as a ratio of ratios to show an estimation of the increase in RT at a specific time (t) compared to time 0, i.e. no depletion (0). The formula used is as follows:

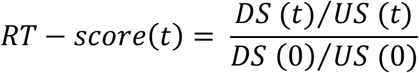

To filter out small and insignificant changes in these values, a cut-off value of 1.1 with an adjusted p-value < 0.05 (based on a t-test with Benjamini-Hochberg FDR corrections) were implemented, based on the distribution of these scores (see Figure 4 and https://github.com/ppolg/Mtb_termseq/blob/main/R/Mtb_perdistance.R) and the distribution of significant TTS/RT scores with varying thresholds calculated (https://github.com/ppolg/Mtb_termseq/blob/main/R/Mtb_pvalues.R; see Figure S6 for relevant heatmaps). These significance thresholds were used to generate the Venn diagrams in Figure 5.

For CondTTS (see below), the 100-nucleotide window was replaced by a specific window for each identified RNA leader, spanning the region from the TSS to the CondTTS, with a same length nucleotide window downstream of the TTS. Using these RNA leader-specific windows, RT-scores and significance were calculated as above.

#### Identification of CondTTS and RNA leaders

TTS localized within a 5’ UTR (defined as region between TSS and annotated ORF) or within the beginning of an annotated ORF (defined within the first quarter of the gene, if a TSS was assigned) were extracted using Bedtools and identified as CondTTS. In case multiple TTS were associated within one region, the strongest signal was identified as CondTTS, and the other signals as SecCondTTS (Secondary CondTTS). Functional gene category enrichment was determined using a hypergeometric test and corrected with a Benjamini-Hochberg multiple testing.

#### uORF identification and re-classification of Orphan TTS

To identify potentially translated upstream ORFs (uORFs), we used Ribo-seq data from Sawyer *et al*.^36^ focusing on RNA leaders. Adapter sequences were trimmed using cutadapt^62^ and reads aligned to the AL123456.3 genome using Bowtie2^56^. Coverage at each nucleotide position was calculated using Bedtools and normalised to counts per million. Ribosome footprints within our identified RNA leaders were extracted using SciPy^63^. To filter out low-confidence peaks, an arbitrary minimum threshold of 250 reads in each biological replicate was applied.

SD-associated uORFs for the windows described above were identified with the find-boxes function from the segmentation-fold tool available on the Galaxy Webserver (https://usegalaxy.eu/), with potential SDs defined as regions of 5 consecutives purines found in RNA leaders. Next, NTG start codons were identified and the distance to the potential SD determined. A uORF was annotated as likely translated if the distance between the 3’ edge of the SD-sequence and the start codon was between 5 and 15 nucleotides (https://github.com/ppolg/Mtb_termseq/blob/main/R/Mtb_uORF_TTS.R).

A similar approach was designed to identify potential leaderless uORFs in RNA leaders associated with an Orphan TTS. ORFs located 5 or less nucleotides downstream of TSS were considered likely leaderless. Ribo-seq peaks within these windows were assigned as leaderless uORFs. Predictions were filtered against already annotated ORFs using Bedtools^58^ and with a minimal ORF length of 5 amino acids.

From the 123 Orphan TTS, a new search was conducted using Bedtools to find TTS localised within uORFs. The Orphan TTS found in those windows were re-assigned as uORF-Internal.

#### Identification of overlapping ORFs in *M. tuberculosis*

Identification of overlapping ORFs was performed using R and based on the latest release of the MycoBrowser annotation (Release 4; 2021-03-23)^54^. The code and corresponding figures are available at https://github.com/ppolg/Mtb_termseq/blob/main/R/Mtb_NUGA.R. Overlapping ORFs were grouped by functional category based on the Mycobrowser annotation. Plots were drawn using ggplot2^64^. Significance of co-occurrence (see Figure 5F) was calculated using Fisher’s exact test with BH correction for FDR.

#### Additional data analysis

Statistical testing and additional data analysis, including differential gene expression analysis for RNA-seq and Term-seq, plotting of RNA-seq depth around TTS based on Term-seq data and generation of subsequent generation of heatmaps was performed in R. The codes used to generate the results and the corresponding figures are available at https://github.com/ppolg/Mtb_termseq. This GitHub repository contains a more detailed description of the specific code used for data analysis and figure generation, and the individual files with code contain detailed annotations. All analysis was based on the H37Rv reference genome FASTA file from NCBI. All plots were drawn using ggplot2 ^64^. Differential expression analysis for RNA-seq and Term-seq data was carried out with DEseq2 ^65^ based on HTseq-count of the BAM files^66^. Calculation of RNA-seq coverage around the TTS (Figure 4D and 4H, https://github.com/ppolg/Mtb_termseq/blob/main/R/Mtb_plotdepth.R) was performed by using counts per million read coverages, normalised to the mean coverage 50-75 nucleotides upstream of the TTS and expressed as a log2-fold change for both aggregate plots and all heatmaps.

### QUANTIFICATION AND STATISTICAL ANALYSIS

The proportion of mono- and tri-phosphorylated TSS as shown in Figure 1B were tested with a hypergeometric test use with R and corrected with a Benjamini-Hochberg (BH) multiple testing correction.

The enrichment of TTS classification as shown in Figure 3C was tested with a Chi-square and corrected with BH.

Enrichment of Internal TTS within the first quarter of annotated ORF, as shown in Figure 3D, was assessed with Student’s t-test.

Significance of overlap between mapped TTS and predicted RDTS, as shown in Supplementary Figure 3A was calculated using hypergeometric test with BH corrections based on 2567 TTS, 29096 RDTS and 38596 RD regions of 228 nucleotides.

Significance of TTS scores and RT scores was assessed with a t-test for each TTS between T0 and each time point, and corrected with BH in R (https://github.com/ppolg/Mtb_termseq/blob/main/R/Mtb_pvalues.R), as shown in Supplementary table 4 and 5.

Overlaps between TTS score and RT score were assessed with a Fisher’s exact test with BH corrections.

Representation of CondTTS in different genes based on ontology, as shown in Supplementary Figure 7, was tested with a hypergeometric calculation, corrected with BH for multiple sampling errors.

The significance of premature termination presence within translated RNA leaders was assessed with a hypergeometric test, corrected with BH.

Prevalence of the different categories of overlapping ORFs with gene ontology categories, as shown in Figure 6E were tested with hypergeometric test and corrected with BH. Co-occurrence of overlaps, as shown in Figure 6F, was tested with a Fisher’s exact test, corrected with BH (https://github.com/ppolg/Mtb_termseq/blob/main/R/Mtb_NUGA.R).

