## Supplementary figures for "Conditional termination of transcription is shaped by Rho and translated uORFS in *Mycobacterium tuberculosis*"

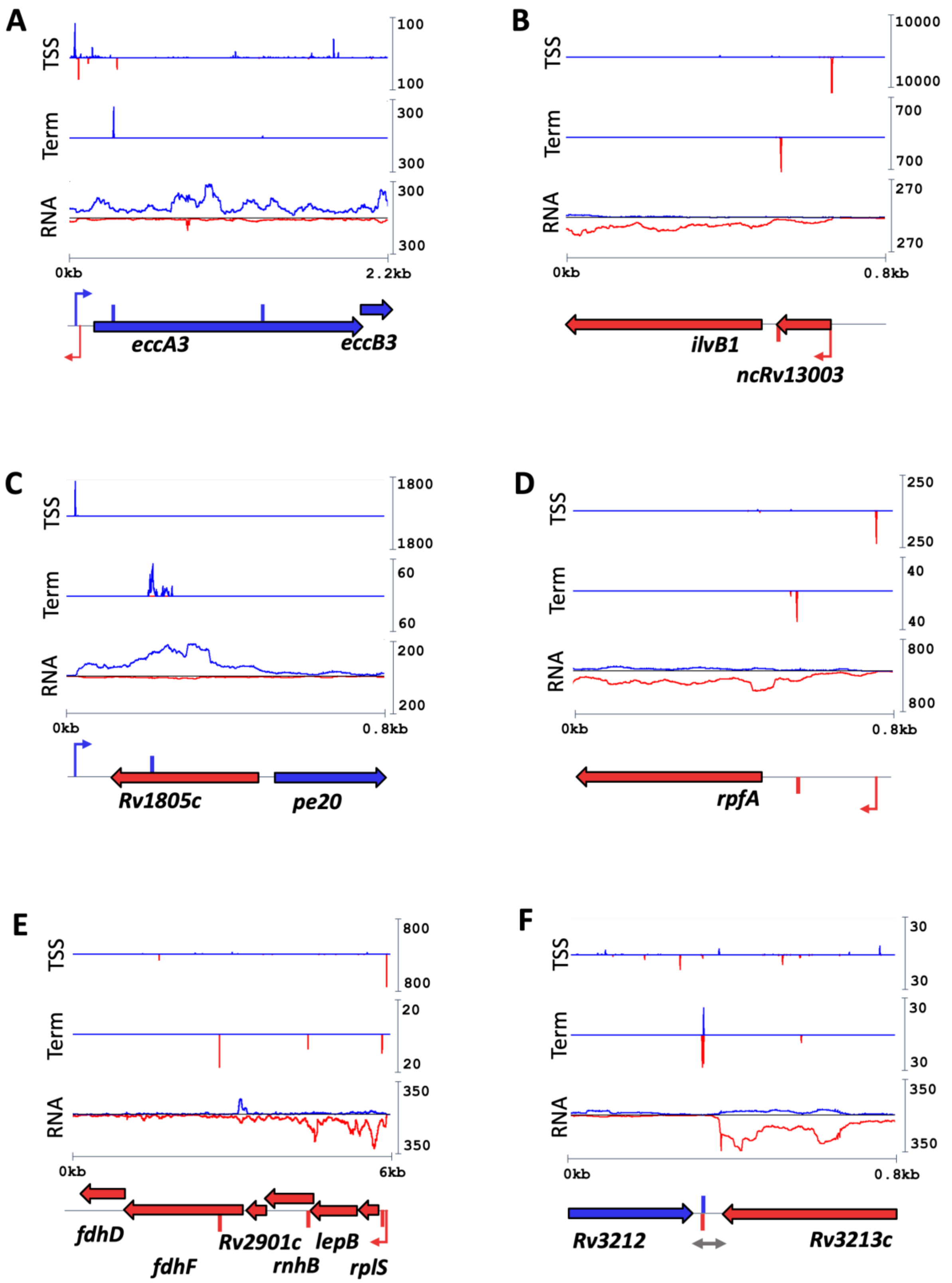


**Supplementary Fig. 1: Additional TTS profiles**. Additional profiles from individual, sharp peaks (A, B); a cluster of peaks within a defined region (C, D), multiple, low-intensity peaks covering entire genes or operons (E, F) and overlapping peaks (G, H). Blue traces: Coverage on the plus strand. Red traces: Coverage on minus strand. Blue / Red arrow: ORF. Blue / Red bar: Mapped TTS. Blue / Red thick arrow: Mapped TSS. Grey arrow: Position of predicted TRIT (Gardner et al., 2011).

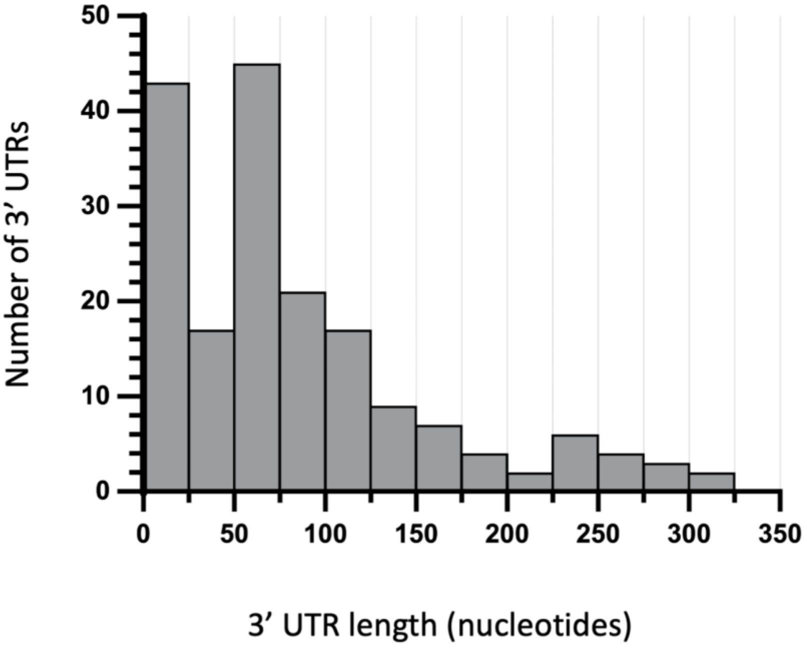


**Supplementary Fig. 2: Distribution of 3’ UTR lengths**. The length of 3’ UTRs in nucleotides was plotted after calculating the distance from the Stop Codon to nearest Final TTS downstream. Grey bars indicates number of 3’ UTRs assigned for each 25-nucleotide window.

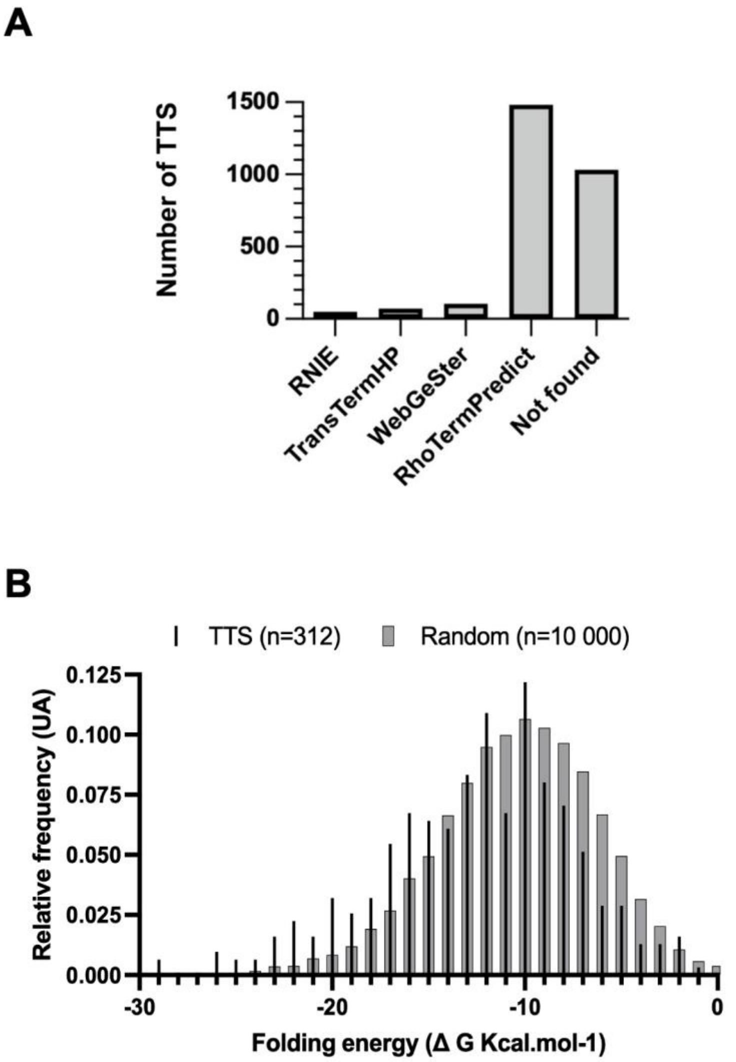


**Supplementary Fig. 3: Predicted mechanism of termination of transcription**. **A**. Experimentally identified TTS were compared to predicted terminators obtained by applying the indicated algorithms to *M. tuberculosis* H37Rv (AL123456.3) genome. TTS were compared to TRITs that have been predicted by RNIE (Gardner et al., 2011), to L-shaped intrinsic terminators for TransTermHP (Kingsford et al., 2007), to L- and I-shaped intrinsic terminators from WebGeSter (Mitra et al., 2011) and to Rho-dependent terminators predicted with RhoTermPredict (RTP; Di Salvo et al., 2019). Hypergeometric test with BH correction indicated that the overlap between mapped TTS and RTP was highly significant (p-value < 0.004. **B** Folding energy around each TTS compared to folding energy around random positions was calculated

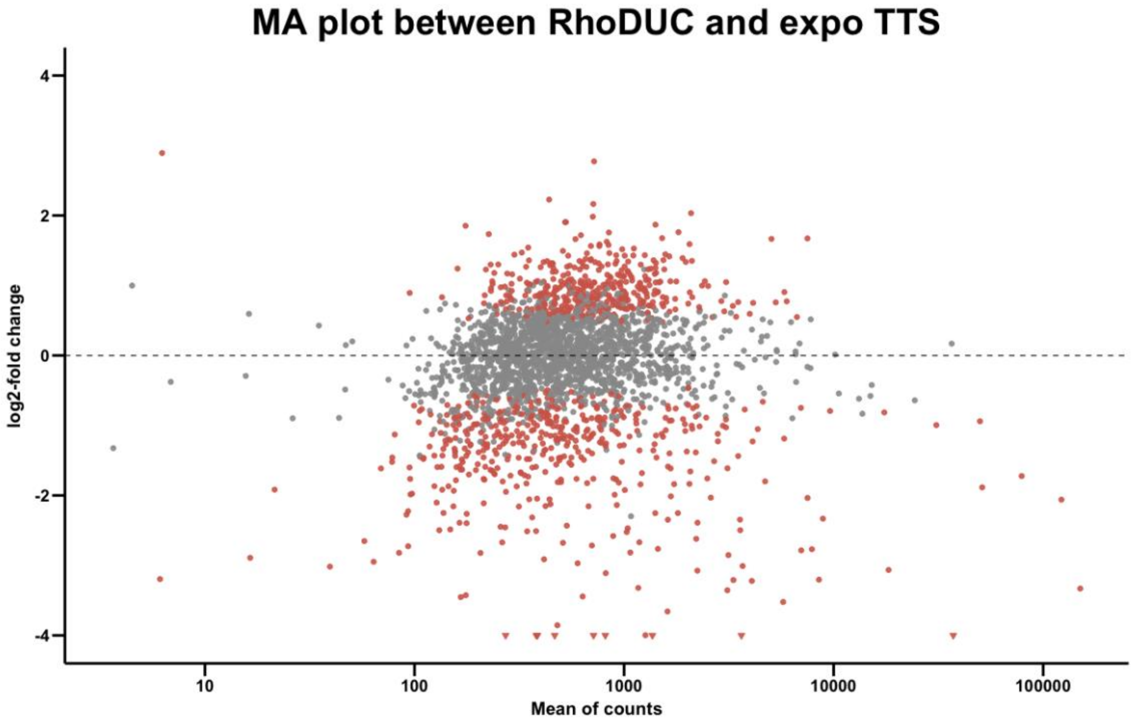


**Supplementary Fig. 4: RhoDUC strain TTS comparison from H37Rv**. (A) MA-plot showing the variation in TTS coverage between H37Rv and RhoDUC strains. The read counts from experimentally identified TTS in H37Rv were extracted from the RhoDUC strain in a window of 5 nucleotides and compared to the number in H37Rv. Red dots show TTS with a $\log_2 \geq 1$ or $\log_2 \leq -1$.

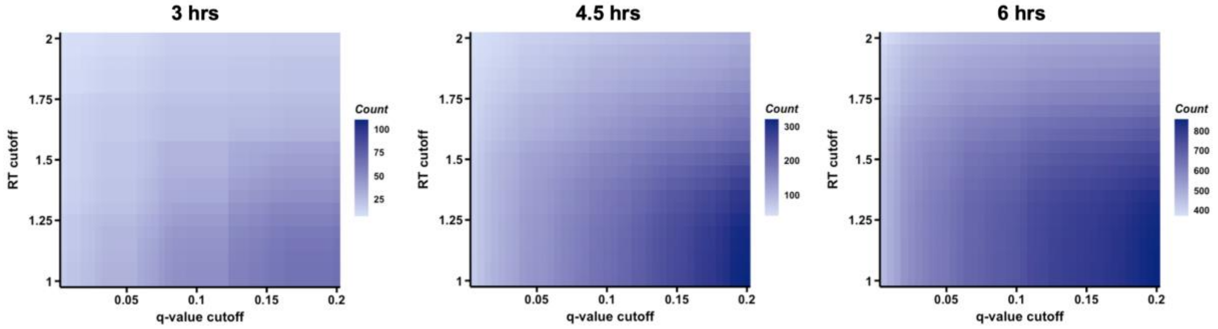


**Supplementary Fig. 5**. Classification of Rho-dependent TTS. The heatmaps show the number of TTS which would be classified as Rho-dependent at the various time points. Horizontal axis indicates the stringency of statistical testing (corrected with Benjamini-Hochberg FDR), vertical axis shows the chosen cutoff for RT scores. Scaling bars indicated on the right show absolute number of CondTTS. Note the different scaling bars.

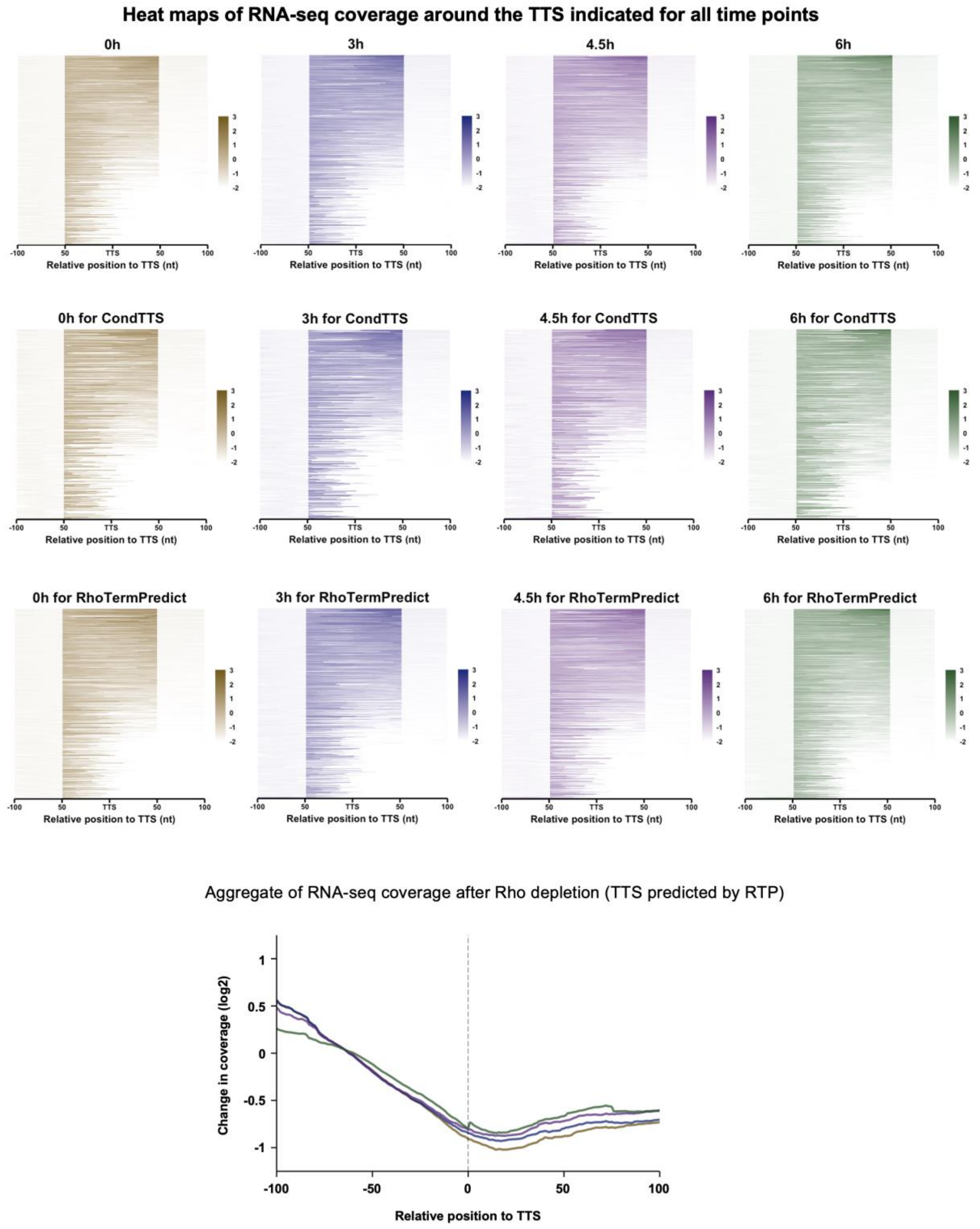


**Supplementary Fig. 6: Heatmaps of RNA-seq coverage around the mapped TTS.** RT-scores were calculated for each time point based on the ratio of normalised read coverage upstream and downstream of each TTS (see methods). Values were plotted in heatmaps for each timepoint for All TTS, for CondTTS and for TTS falling in predicted Rho-dependent regions. Below, aggregate plot summarising heatmaps from TTS predicted by RhoTermPredict (RTP) (Di Salvo et al., 2019)

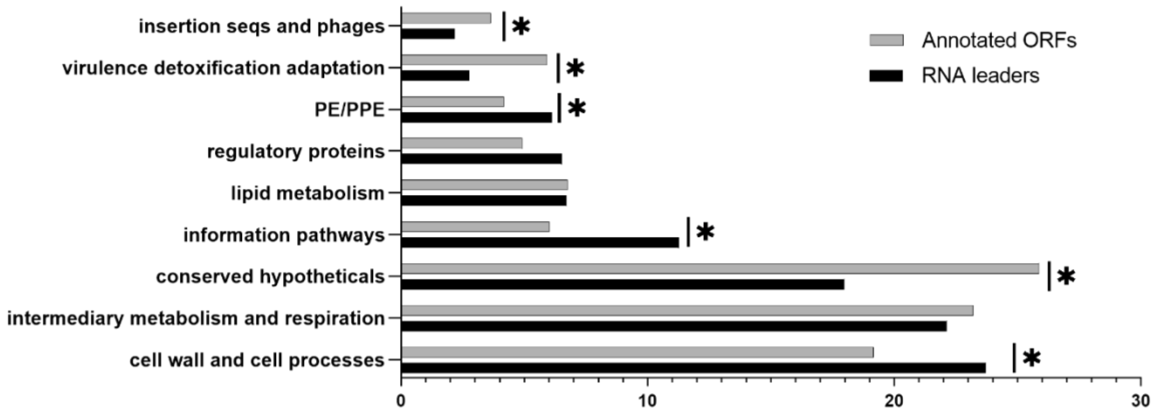


**Supplementary Fig. 7: Distribution of RNA leaders according to functional gene category**. RNA leaders were classified according to the functional category from Mycobrowser of the annotated ORF they are assigned. Enrichment was validated with hypergeometrical tests (with Benjamini-Hochberg correction for FDR) for each category with a p-value ≤ 0.05 (Star). Black bars: Proportion in % of RNA leaders. Grey bars: Proportion in % of annotated ORF.
