## Supplementary Table 8 for "Conditional termination of transcription is shaped by Rho and translated uORFS in *Mycobacterium tuberculosis*"

**Supplementary table 9: strains & plasmids**

| **Strains and plasmids** | **Description** | **Reference** |
| --- | --- | --- |
| **Strains** |  |  |
| *M. tuberculosis* |  |  |
| H37Rv | Virulent model organism of *M. tuberculosis* | [1] |
| Rho-DUC | Δ*rho*::*hygR*, att-L5::rev*tetR*-P^tetOFF^-*rho*DAS-*zeoR*, Tweety::*tetR*-P^tetOFF^-*sspB*-*kanR.* Rho-inducible deletion strain with anhydrous tetracyclin. | [2] |
| *M. smegmatis* |  |  |
| MC^2^ 155 | High-frequency transformation mutant of *M. smegmatis* | [3] |
| *E. coli* |  |  |
| DH5α | Cloning strain for plasmids construction | NEB C2987 |
| **Plasmids** |  |  |
| pIRaTE | Translation fusion reporter plasmid carrying the rrnB promoter sequence and lacZ gene downstream | [4] |
| pIRaTE2020 | pIRaTE derivative ith the insertion of the pIRaTE2020 insert between XhoI and HindIII sites, carrying a linker, the *rrnB* and PCL1 core promoters in opposite directions and an intrinsic terminator in between | This work |
| pIRaTE2020_urpfA | pIRaTE2020 with the Fusion_uRpfA DNA fragment inserted between HindIII and NcoI sites | This work |
| pIRaTE2020_urne | pIRaTE2020 with the Fusion_uRne DNA fragment inserted between HindIII and NcoI sites | This work |
| pIRaTE2020_u1glyA2 | pIRaTE2020 with the Fusion_u1GlyA2 DNA fragment inserted between HindIII and NcoI sites | This work |
| pIRaTE2020_uilvB1 | pIRaTE2020 with the Fusion_uIlvB1 DNA fragment inserted between HindIII and NcoI sites | This work |
| pIRaTE2020_udnaA | pIRaTE2020 with the Fusion_uDnaA DNA fragment inserted between HindIII and NcoI sites | This work |
| pIRaTE2020_u2pe20 | pIRaTE2020 with the Fusion_u2Pe20 DNA fragment inserted between HindIII and NcoI sites | This work |
| pIRaTE2020_u2Rv1535 | pIRaTE2020 with the Fusion_u2Rv1535 DNA fragment inserted between HindIII and NcoI sites | This work |
| pIRaTE2020_NoSD | pIRaTE2020 with the Fusion_NoSD DNA fragment inserted between HindIII and NcoI sites | This work |

1.	Cole, S.T., et al., *Deciphering the biology of Mycobacterium tuberculosis from the complete genome sequence.* Nature, 1998. **393**(6685): p. 537-44.

2.	Botella, L., et al., *Depleting Mycobacterium tuberculosis of the transcription termination factor Rho causes pervasive transcription and rapid death.* Nat Commun, 2017. **8**: p. 14731.

3.	Snapper, S.B., et al., *Isolation and characterization of efficient plasmid transformation mutants of Mycobacterium smegmatis.* Mol Microbiol, 1990. **4**(11): p. 1911-9.

4.	Moores, A., et al., *Expression, maturation and turnover of DrrS, an unusually stable, DosR regulated small RNA in Mycobacterium tuberculosis.* PLoS One, 2017. **12**(3): p. e0174079.
