## Supplementary Table 9 for "Conditional termination of transcription is shaped by Rho and translated uORFS in *Mycobacterium tuberculosis*"

**Supplementary table 9: primers, oligos, gBlock**

| **DNA name** | **Sequence** | **Experiment / Manipulation** |
| --- | --- | --- |
| **Reporter primers** |  |  |
| Fusion_uRpfA_F | AGCTTCCGAACTCCATCCAGTGGGCGAGGGATTCCGACCGCGGATGGAGGCGGGGGACGC | Oligo Annealing and ligation (HindIII and NcoI sites) |
| Fusion_uRpfA_R | CATGGCGTCCCCCGCCTCCATCCGCGGTCGGAATCCCTCGCCCACTGGATGGAGTTCGGA | Oligo Annealing and ligation (HindIII and NcoI sites) |
| Fusion_u1GlyA2_F | AGCTTCGAACAGCCGCTGACGCGATGTGGGAGAACCTCCATGTCGAGGCGCCGTGC | Oligo Annealing and ligation (HindIII and NcoI sites) |
| Fusion_u1GlyA2_R | CATGGCACGGCGCCTCGACATGGAGGTTCTCCCACATCGCGTCAGCGGCTGTTCGA | Oligo Annealing and ligation (HindIII and NcoI sites) |
| Fusion_uRne_F | GCCGGATTTGTATTAGACTAAGCTTGGGTGAACGTCGCCGAAGCGGTGTCGGACGCCCGAATGAAGACCATGGATGATCCCGTCGTTTTACA | Oligo Annealing and Gibson Assembly |
| Fusion_uRne_R | TGTAAAACGACGGGATCATCCATGGTCTTCATTCGGGCGTCCGACACCGCTTCGGCGACGTTCACCCAAGCTTAGTCTAATACAAATCCGGC | Oligo Annealing and Gibson Assembly |
| Fusion_uDnaA_F | GCCGGATTTGTATTAGACTAAGCTTCGGGTGTTTTCAACACGAGGATCGCGAGCCGTTGCCGGTAGGTTGCGCATGGATGATCCCGTCGTTTTACA | Oligo Annealing and Gibson Assembly |
| Fusion_uDnaA_R | TGTAAAACGACGGGATCATCCATGCGCAACCTACCGGCAACGGCTCGCGATCCTCGTGTTGAAAACACCCGAAGCTTAGTCTAATACAAATCCGGC | Oligo Annealing and Gibson Assembly |
| Fusion_u2Pe20_F | ATTTGTATTAGACTAAGCTTCTGTGCATCGGCATCCCCGTGTGCCCCGGCCGTGAGGAGGTGAGAGCGAAATGAGTCCCCCCATGGATGATCCCGTCGTT | Oligo Annealing and Gibson Assembly |
| Fusion_u2Pe20_R | AACGACGGGATCATCCATGGGGGGACTCATTTCGCTCTCACCTCCTCACGGCCGGGGCACACGGGGATGCCGATGCACAGAAGCTTAGTCTAATACAAAT | Oligo Annealing and Gibson Assembly |
| Fusion_u2Rv1535_F | ATTTGTATTAGACTAAGCTTTATCGTGCGGACAACCGTACGTGTCGTGGCCGTGAGGAGGTGAGGGACGCATGAGTTCCCCCATGGATGATCCCGTCGTT | Oligo Annealing and Gibson Assembly |
| Fusion_u2Rv1535_R | AACGACGGGATCATCCATGGGGGAACTCATGCGTCCCTCACCTCCTCACGGCCACGACACGTACGGTTGTCCGCACGATAAAGCTTAGTCTAATACAAAT | Oligo Annealing and Gibson Assembly |
| Fusion_uIlvB1_F | GCCGGATTTGTATTAGACTAAGCTTCTATGGACAAGGCCGGAAAGCCCGGGATGCTCGTAGTAATTGGCATGGATGATCCCGTCGTTTTACA | Oligo Annealing and Gibson Assembly |
| Fusion_uIlvB1_R | TGTAAAACGACGGGATCATCCATGCCAATTACTACGAGCATCCCGGGCTTTCCGGCCTTGTCCATAGAAGCTTAGTCTAATACAAATCCGGC | Oligo Annealing and Gibson Assembly |
| Fusion_NoSD_F | GCCGGATTTGTATTAGACTAAGCTTGAGTAGGAGATTTTCACCTCCTTTCCTTCCTACCATGGATGATCCCGTCGTTTTACA | Oligo Annealing and Gibson Assembly |
| Fusion_NoSD_R | TGTAAAACGACGGGATCATCCATGGTAGGAAGGAAAGGAGGTGAAAATCTCCTACTCAAGCTTAGTCTAATACAAATCCGGC | Oligo Annealing and Gibson Assembly |
| **PCR primers** |  |  |
| RpfB-KsgA_RT_F2 | AACGGCGGGCTGCGGTATGC | PCR on total RNA (gDNA contaminants) |
| RpfB-KsgA_RT_R2 | CGCACCGTGTTGGCGTCGTG | PCR on total RNA (gDNA contaminants) |
| pIR_F | TTGACTCCATTGCCGGAT | PCR to amplify the mutated inserts from PCR2.0 |
| pIR_R | GACGTTGTAAAACGACGGGA | PCR to amplify the mutated inserts from PCR2.0 |

gBlock for changing region between promoters in pIRaTE

ctcgagTACGTATTAATTAAactagtAAGTTAcgtccttggaaactgggAGTCAAatccccaggtcagagggctattttccctggtcagacgcggtcacccTTTAAAagacaaatccgccgagcttcgacGCCCGCCTAATGAGCGGGCTTTTTTTTtcctgcaggattctagacgacTTGACTccattgccggatttgtatTAGACTaagctt
